# Male gut microbiome mediates post-mating sexual selection in *Drosophila*

**DOI:** 10.1101/2025.05.23.655737

**Authors:** Komal Maggu, Abhishek Meena, Alessio N. De Nardo, Sonja H. Sbilordo, Jeannine Roy, Stefan Lüpold

**Author notes:** Komal Maggu & Stefan Lüpold, **Email:**.

## Abstract

When females mate with multiple males, sexual selection continues after mating and favors males with more competitive ejaculates. While these traits are physiologically costly and sensitive to internal state, the effects of the gut microbiome on male reproductive performance remains underexplored. Here, we tested how gut microbial communities influence sperm competition outcomes and the traits underpinning them by inoculating male *Drosophila melanogaster* with fully factorial synthetic consortia of five common commensal bacteria. We found highly context-dependent and non-additive microbial effects, along with two general patterns: *Acetobacter*-dominated microbiomes enhanced sperm transfer and increased paternity shares by 20–27% relative to germ-free males, whereas species richness strongly predicted sperm viability and female sperm storage. These results indicate that gut microbes contribute to multiple components of ejaculate performance through distinct pathways, potentially including nutrient provisioning for spermatogenesis, acetate-mediated energy metabolism, and microbe-dependent modulation of female sperm use. By integrating microbial ecology with reproductive biology, our findings establish the gut microbiome as an overlooked but fundamental modulator of sexually selected traits. This perspective helps clarify how non-genetic and community-based factors generate phenotypic variation under strong selection and provides a framework for future work on the mechanisms, transmission routes, and evolutionary consequences of host–microbe interactions in reproduction.

## Introduction

In polyandrous species, in which females mate with multiple males, post-mating sexual selection favors males with more competitive ejaculates, such as comprising more and higher-quality sperm or seminal fluid (Simmons & Fitzpatrick, 2012; Fitzpatrick & Lüpold, 2014). Producing such ejaculates is energetically costly and diverts resources from other fitness-related traits such as growth, maintenance, or immunity (Dewsbury, 1982; Olsson *et al*., 1997; Perry & Tse, 2013; Lüpold *et al*., 2016; Macartney *et al*., 2019), which increases the susceptibility of ejaculate traits to internal physiological and external ecological impacts and so can affect male reproductive strategies (Wedell *et al*., 2002; Parker & Pizzari, 2010; Simmons & Fitzpatrick, 2012).

A commonly invoked framework to explain variation in costly traits is condition dependence, where trait expression reflects an individual’s ability to acquire and allocate resources under physiological or environmental constraints (Rowe & Houle, 1996; Cotton *et al*., 2004; Tomkins *et al*., 2004). Traditionally, variation in condition is assumed to be driven primarily by genetic variation in mechanisms such as nutrient assimilation, metabolic regulation, or stress resilience (Hill, 2011). However, this view may overlook the host’s microbial symbionts as another critical source of phenotypic variation. In particular, the gut microbiome has emerged as a powerful regulator of host physiology by influencing nutrient acquisition (Douglas, 2009; Shin *et al*., 2011; Alcock *et al*., 2014; Fetissov, 2017), resource allocation (Emelianoff *et al*., 2008; Gould *et al*., 2018; Walters *et al*., 2020), or hormone signaling (Qi *et al*., 2021), with likely consequences for reproductive performance (Cai *et al*., 2022).

As a dynamic and metabolically active community, the microbiome alters the physiological landscape in which reproductive traits are developed and expressed (Lv *et al*., 2024). Consequently, microbial communities may modulate—or even drive—variation in ejaculate traits traditionally ascribed to reflect intrinsic male quality (Lv *et al*., 2024). These effects often arise independently of host genetic condition (Spor *et al*., 2011; Koskella *et al*., 2017) and may thus establish an ecologically responsive axis of phenotypic variation under sexual selection. Unlike classic determinants of condition like diet or infection (Hill, 2011), the microbiome operates as a functionally integrated, host-associated system that is reciprocally shaped by host genotype, behavior, and ecology while actively modulating host physiology (Archie & Theis, 2011; Foster *et al*., 2017; Moeller *et al*., 2017). This reciprocal relationship persists even in systems with transient microbial colonization, such as *Drosophila*, where microbiome stability *is influenced by host* diet (Blum *et al*., 2013; Pais *et al*., 2018). Moreover, vertical, social, or sexual transmission of the microbiota enables co-regulation and evolutionary feedback unlike other environmental effects (Robinson *et al*., 2019). Microbial communities may thus act as dynamic contributors to reproductive phenotypes through host interactions, and ultimately to the evolution of sexually selected traits (Rosenberg & Zilber-Rosenberg, 2018; Rowe *et al*., 2020). While such microbiome-mediated modulation in female reproduction has been widely studied, including effects on egg production and mate choice in *Drosophila* and other taxa (Sharon *et al*., 2010; Gould *et al*., 2018; Comizzoli *et al*., 2021; Chadchan *et al*., 2022), its role in males remains remarkably underexplored, even more so in the context of sexual selection (Rowe *et al*., 2020). Existing research has largely focused on humans, rodents, and livestock, and has primarily examined broad associations between dysbiosis and impaired testicular function (Al-Asmakh *et al*., 2014; Ding *et al*., 2020) or the influence of the seminal microbiome on *in vitro* sperm parameters in a clinical or agricultural context (Altmäe *et al*., 2019; Rowe *et al*., 2020; Farahani *et al*., 2021). However, the role of the gut microbiota in influencing the outcome of competitive fertilization, particularly through condition-dependent ejaculate traits and female-mediated processes like sperm retention and utilization, remains untested. This gap is especially surprising given the near-universal presence of post-mating reproductive processes, such as sperm competition and differential female sperm handling, that shape both fertilization success and the evolution of reproductive traits across animals (Simmons & Fitzpatrick, 2012; Firman *et al*., 2017; Lüpold & Pitnick, 2018).

Here, we tested the hypothesis that the gut microbiome modulates male reproductive fitness under competitive fertilization by enhancing condition-dependent ejaculate traits in *Drosophila melanogaster*, a model for tractable host-microbe interactions (Ridley *et al*., 2013; Wong *et al*., 2013; Douglas, 2018). To this end, we inoculated germ-free (axenic) males with all 31 non-axenic combinations of five gut bacteria (*Acetobacter orientalis*, *A. tropicalis*, *A. pasteurianus*, *Lactobacillus brevis*, and *L. plantarum*), while maintaining an axenic control group. These bacterial species have well-characterized metabolic and reproductive effects in female *D. melanogaster* and contrasting functional profiles that may differentially affect host physiology (Gould *et al*., 2018; Matthews *et al*., 2021). Given the shared physiological pathways between sexes in this species, these bacteria can be predicted to also impart potentially critical effects on male fitness. Further, when present together, they can recapitulate key traits of intact microbiomes (Supplementary Fig. S1). We then quantified competitive fertilization success (i.e., second male’s paternity share, P₂) and its mechanistic underpinnings in our experimental males: sperm viability, male sperm transfer, and female sperm retention and storage in response to different microbiome parameters (Fig. 1A).

**Figure 1.**
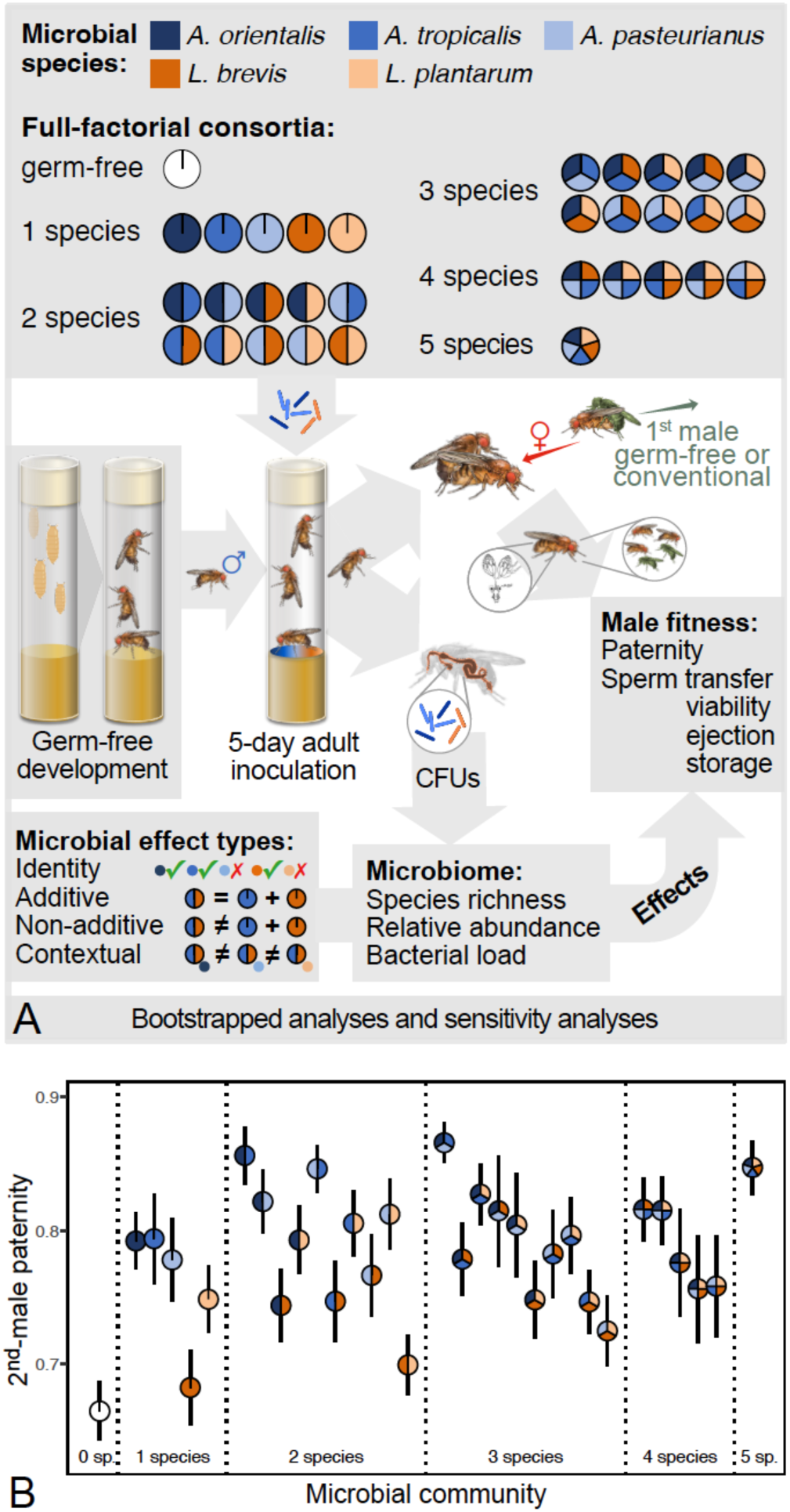
Visual representation of the experimental workflow (A) and variation in focal-male sperm competitiveness relative to their microbial community (B). In panel A, experimental males either remained germ-free or were inoculated with one of the 31 full-factorial combinations of *Acetobacter* and *Lactobacillus* species (indicated by shades of blue and orange, respectively). Five days later, they inseminated a previously mated female (germ-free, first-mated with a germ-free or a conventional male) before we counted offspring for paternity shares (based on ubiquitously GFP-labeled first males) and dissected either females for sperm counts (ejected and retained) or males for sperm viability. In parallel, males were analyzed for microbial identity and abundance (based on colony-forming units, CFUs). Microbiome and reproductive data were then combined across 1,000 bootstrapped samples for formal analyses and effect validation through sensitivity analyses. Error bars in panel B reflect 95% confidence intervals from raw distributions (*N* = 20.2 ± 2.95 (± s.d.) trials per consortium).

We predicted that (i) males with diverse microbiomes outperform germ-free or low-diversity males due to improved nutrient acquisition (Douglas, 2009; McMullen *et al*., 2020), (ii), competitive advantages are shaped by bacterial identity, including species with known metabolic contributions to reproductive function (e.g., *Acetobacter* spp.; Gould *et al*., 2018), as well as by synergistic interactions and emergent properties of community composition (e.g., *Acetobacter* dominance (Gould *et al*., 2018)), and (iii) microbiome-driven gains in sperm viability, transfer efficiency and competitive storage underpin fertilization success (Lüpold *et al*., 2012, 2020; Tourmente *et al*., 2019). By demonstrating that gut microbiome composition and diversity modulate key ejaculate traits and competitive fertilization success, this study bridges microbial ecology and sexual selection theory, highlighting an underappreciated source of phenotypic variation in traits under strong post-mating selection.

## Results

### Microbial effects on competitive fertilization success

To test if gut microbes modulate male reproductive fitness under post-mating sexual selection, we evaluated competitive fertilization success in male *D. melanogaster* inoculated with all 32 combinations of five gut bacterial species (mentioned above). This full-factorial approach disentangled species- and community-level effects using validated gnotobiotic methods (*Material and Methods*; Supplementary Figs. S1 and S2). Males competed in sequential matings against either germ-free or conventional rivals (with unmanipulated microbiomes), which provided distinct competitive contexts to isolate intrinsic microbiome benefits. Competitor males, which mated with standardized axenic females three days before focal males, ubiquitously expressed green fluorescent protein (GFP) for paternity assignment (Lüpold *et al*., 2012, 2013).

Across both competitive contexts, fertilization success increased with microbiome diversity. Against axenic rivals, second-male paternity (P_2_) increased with species richness (generalized linear mixed models (GLMM) repeated across 1,000 bootstrapped datasets, accounting for block, microbial combination and overdispersion: *N* = 646 across 32 microbial combinations: odds ratio, *OR* = 1.09 [95% confidence limits: 1.00, 1.15]). Males with all five species gained 27% in P_2_ relative to axenic males (mean P_2_ ± s.d. = 0.85 ± 0.05 vs. 0.67 ± 0.05, respectively; Fig. 1A). Similar gains occurred against conventional rivals (*N* = 624, *OR* = 1.05 [1.00, 1.10]; 20% gain for five-species consortia: 0.72 ± 0.08 vs. 0.60 ± 0.03), confirming that microbiome-driven benefits stem from intrinsic physiological enhancement, not reduced competitor performance.

To investigate microbial community composition in non-axenic males, we analyzed the proportion of *Acetobacter* species (based on colony-forming unit [CFU] counts), species richness, and total bacterial load (log CFUs) using bootstrapped generalized linear models (GLMs) on standardized predictors (*Materials and Methods*). To address moderate collinearity between species richness and bacterial load (variance inflation factor ≈ 6.6), we performed sensitivity analyses including residualization, data subsetting, variable exclusion, ridge regression, variance partitioning, and simulation-based validation (Supplementary Text, Tables S1–S10, Fig. S3), which confirmed the robustness of our estimates. In competitions against axenic control males, *Acetobacter*-dominated microbiotas strongly enhanced paternity success (P_2_: *OR* = 1.21 [1.18, 1.25]), whereas species richness and bacterial load showed negligible effects (*OR* = 1.02 [0.94, 1.11] and 1.03 [0.94, 1.13], respectively; Fig. 2). These patterns held against conventional rivals (proportion *Acetobacter*: *OR* = 1.10 [1.08, 1.13]; richness: *OR* = 1.02 [0.95, 1.08]; load: *OR* = 1.03 [0.96, 1.09]). Variance partitioning confirmed the dominant role of *Acetobacter* by attributing 70–93% of the explainable variance in P_2_ to the unique effect of relative *Acetobacter* abundance, with negligible unique contributions from richness or load (Supplementary Tables S7 and S8; Supplementary Fig. S4).

**Figure 2.**
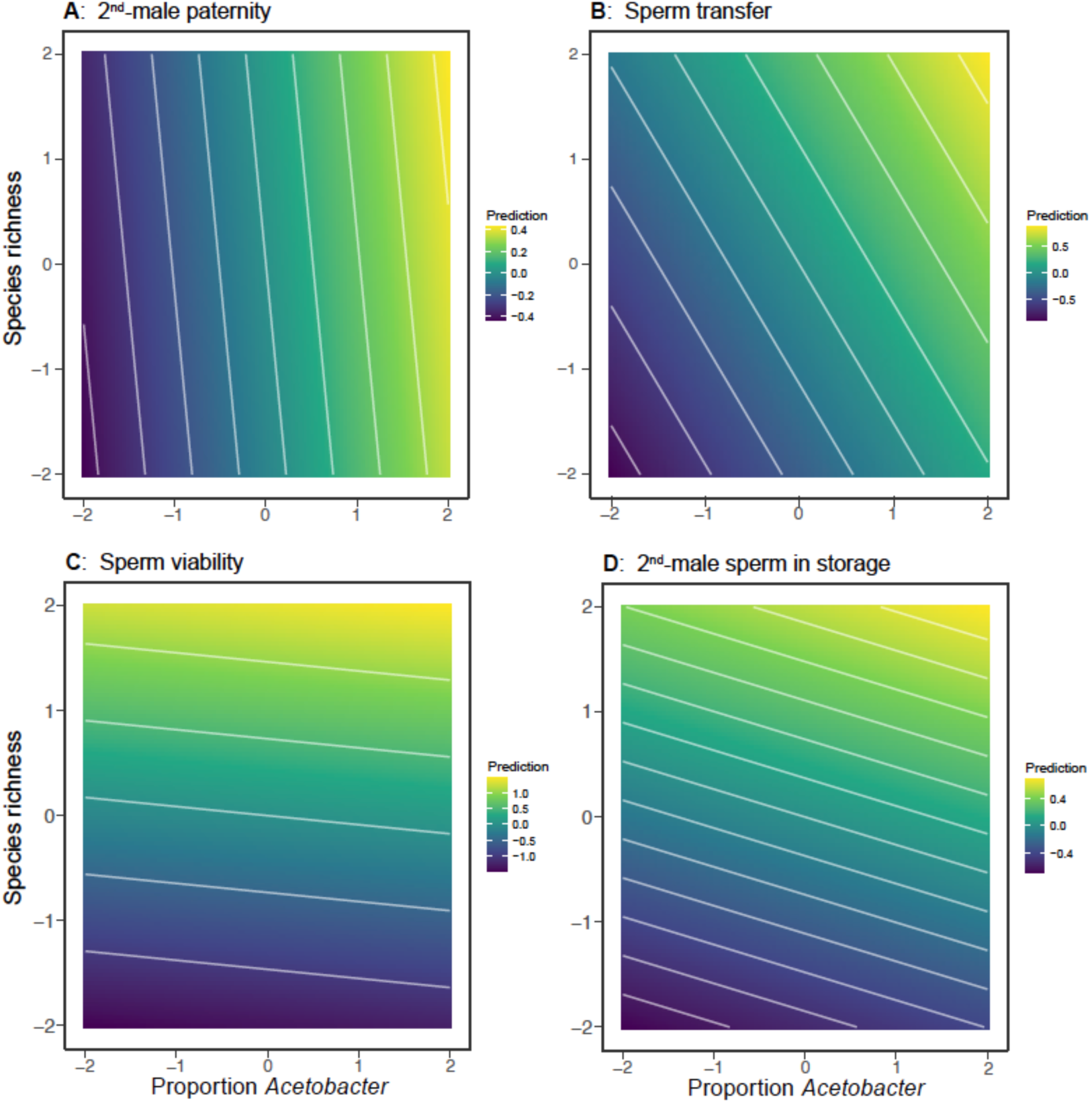
Effects of the total number of bacterial species and the proportion of *Acetobacter* representation in the microbiota on the different fitness traits after controlling for total bacterial load. All predictor variables were standardized before analysis, and their response predictions were retrieved from bootstrapped (generalized) linear models (*N* = 1,000 iterations). The axenic treatment was excluded due to undefined microbe abundances. The time to female sperm ejection is omitted here as neither predictor had a significant effect.

Next, we tested for additive microbial effects, using bootstrapped generalized linear mixed models (GLMMs) on *P_2_*, controlling for experimental block, as these analyses involved only response variables with consistent block structures. Five-way interactions were not robust (axenic rivals: *N* = 646, *OR* = 0.82 [0.55, 2.66]; conventional rivals: *N* = 624, *OR* = 0.76 [0.40, 1.39]), but excluding them revealed a four-way interaction of *At* × *Ap* × *Lb* × *Lp* against axenic rivals (*OR* = 1.84 [1.22, 2.77]) and *Ao* × *Ap* × *Lb* × *Lp* against conventional rivals (*OR* = 0.67 [0.49, 0.93]). Numerous two- and three-way interactions further confirmed this non-additivity (Supplementary Figs. S5 and S6).

We then used predictive modeling to estimate reproductive performance in each multi-species consortium based on average values from the corresponding mono-association treatments (e.g., mean of *At*, *Ap*, and *Lb* to predict the *At*-*Ap*-*Lb* combination). These single-species averages accurately predicted only 8 out of 26 (30.8%) multi-species effects (Fig. 3A) against axenic rivals, and 34.6% (9/26) against conventional males. Pairwise averages (e.g., *At*-*Ap*, *At*-*Lb* and *Ap*-*Lb* to predict *At*-*Ap*-*Lb*) were similarly inaccurate (31.2% (5/16) and 37.5% (6/16), respectively; Fig. 1B), highlighting the emergent properties of more complex microbial consortia.

**Figure 3.**
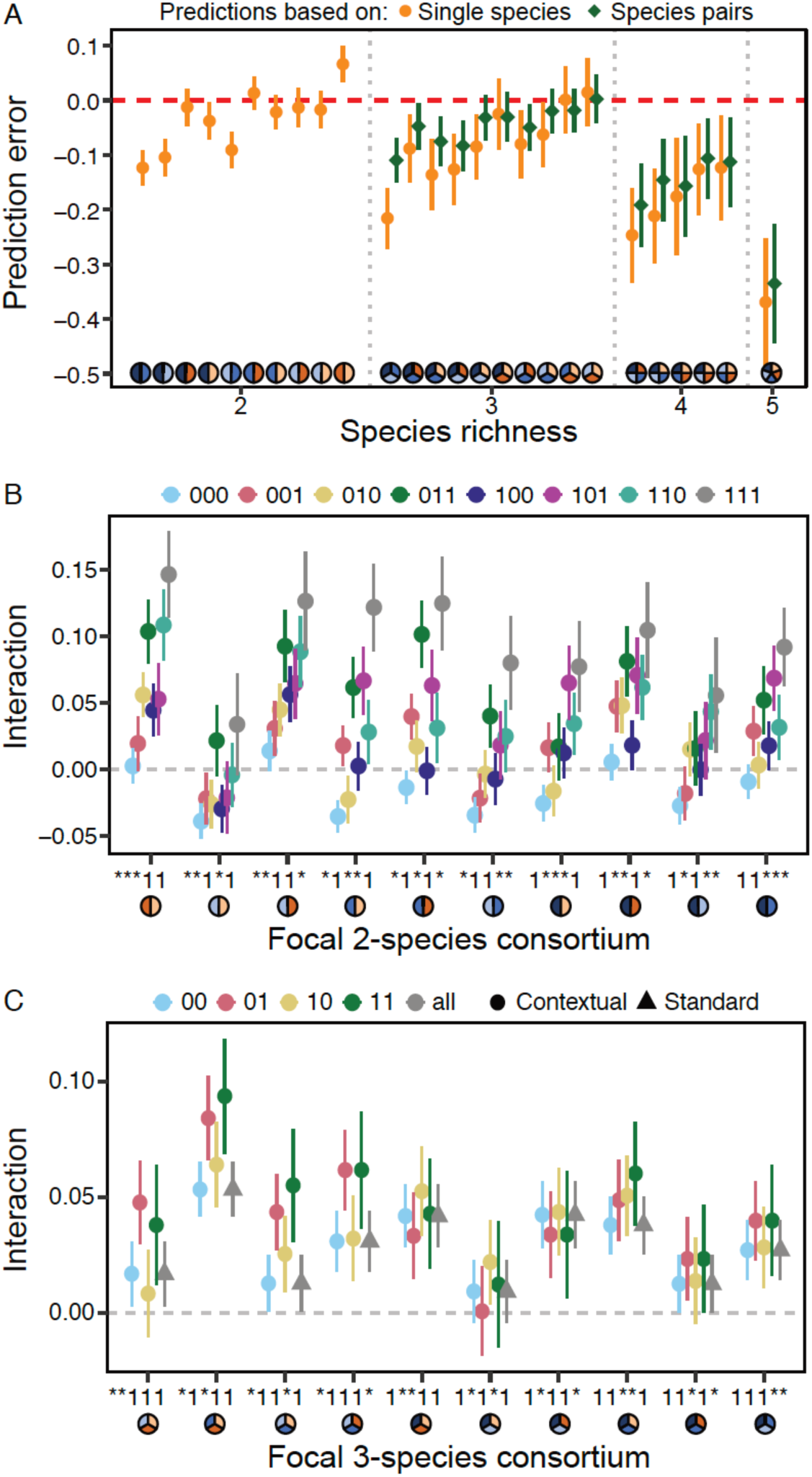
Microbiome effects on the proportion of focal-male paternity based on additive *versus* non-additive interactions among species of the microbiome itself (A) and in response to bystander species influencing these interactions (B and C). Panel A depicts the deviations from observed trait values when averaging those of the individual species (orange points) or species pairs (green diamonds) that jointly represented the same species communities (all coefficients derived from generalized linear mixed-effects models). Negative values indicate the observed trait value was higher than that predicted by averaging the effects of its component species (i.e., synergy), while a positive value indicates it was weaker (i.e., antagonism). The dashed line at zero represents a perfectly additive effect. The *Acetobacter* and *Lactobacillus* species in each focal microbiome are indicated along the *x*-axis (same colors as in Fig. 1), and error bars reflect 95% confidence intervals from bootstrapped prediction differentials. Panels B and C present interactive effects between bacterial species contingent on bystander species, with two-way interactions between pairs of focal of species (B) and three-way interactions between trios of focal species (C), respectively. For each code along the *x*-axes, 1’s indicate the species configuration of the interaction (also visualized by pies underneath) and asterisks denote placeholders for bystander species (order within codes from left to right: *Acetobacter orientalis*, *A. tropicalis*, *A. pasteurianus*, *Lactobacillus brevis*, *L. plantarum*). The binary codes in the legend above each panel indicate the configuration of bystanders in the order, in which they would fill the placeholders (*) in each focal species configuration (1 = presence, 0 = absence). Panel C further distinguishes between the “standard” tests (grey triangles; i.e., global interaction across all community members present) and “contextual” tests (colored points; i.e., separate interactions between a focal species pair or trio for each bystander combination) per focal trio. All coefficients and propagated standard errors were derived from (generalized) linear mixed-effects models across 1,000 bootstrapped datasets.

Finally, the magnitude and often also the direction (synergism or antagonism) of microbial interactions were context-dependent, with pair and triplet effects varying based on bystander species in the community. For example, the same microbial pair could act synergistically or antagonistically depending on which other species were present (Fig. 3B). This dependence on bystanders also held for three-way interactions between focal species (Fig. 3C), again indicating that microbial communities modulate host phenotypes as integrated systems, beyond additive effects.

Together, these results show how microbiome composition, particularly *Acetobacter* dominance, mediates reproductive success.

### Ejaculate traits and female responses mediate microbiome-driven reproductive gains

To elucidate how microbiome composition influences competitive fertilization success, we examined the number of sperm transferred, proportion of viable sperm, time from mating to female ejection of excess second-male and displaced first-male sperm, and the proportional representation of second-male sperm among all sperm stored by female after ejection (Lüpold *et al*., 2012, 2013, 2020; Tourmente *et al*., 2019). We focused on matings with axenic first males to eliminate rival microbial effects, ensuring clear inference of community composition impacts. For visualization of sperm, all assays except sperm viability involved standard and experimental males that, respectively, expressed green or red fluorescent protein in sperm heads (Manier *et al*., 2010).

Microbiome diversity significantly modulated ejaculate and female sperm-storage dynamics, with mean trait values of multi-associated males exceeding those of axenic or mono-associated males by up to 75% (Supplementary Fig. S7). Using bootstrapped GLMs and linear models (LMs), we jointly analyzed species richness, the proportion of *Acetobacter* in consortia, and bacterial load across the non-axenic males (*Materials and Methods*). Sensitivity analyses again confirmed the robustness of our estimates (Supplementary Tables S1–S10, Figs. S3, S4 and S8).

In these analyses, the influence of microbiome parameters differed across traits, revealing the mechanistic pathways to competitive success. For example, species richness strongly enhanced sperm viability (*OR* = 1.98 [1.75, 2.22]) and the relative number of sperm in storage after female sperm ejection (*OR* = 1.31 [1.21, 1.42]), with its unique effects accounting for 51–56% of their total variance (Supplementary Table S4), but it had no clear effect on sperm transfer (*r* = 0.09 [-0.15, 0.32]) or the timing of female sperm ejection (*r* = 0.03 [-0.22, 0.28]; Fig. 2). By contrast, the proportion of *Acetobacter* in consortia was a consistent positive predictor across most traits (sperm viability: *OR* = 1.06 [1.02, 1.10]; sperm transfer: *r* = 0.33 [0.14, 0.52]; sperm storage: *OR* = 1.08 [1.05, 1.10]), except for the timing of female sperm ejection (*r* = 0.17 [–0.07, 0.41]; Fig. 2). The effects of bacterial load were weaker (sperm transfer: *r* = 0.24 [0.01, 0.46]; sperm storage: *OR* = 1.06 [0.99, 1.13]; ejection timing: *r* = 0.06 [–0.17, 0.31]) or negative (sperm viability: *OR* = 0.80 [0.72, 0.89]), and ridge regression suggested that its estimated influence was unstable due to collinearity with species richness (Supplementary Table S6). While sensitivity analyses confirmed the dominant role of *Acetobacter* and species richness, respectively (Supplementary Tables S1–S10), future studies should manipulate bacterial load independently to fully disentangle density from composition. Finally, consistent with prior studies (Lüpold *et al*., 2012, 2013, 2020; Tourmente *et al*., 2019), all these reproductive traits covaried positively with paternity outcomes, if only weakly so for the timing of female sperm ejection (Fig. 4).

**Figure 4.**
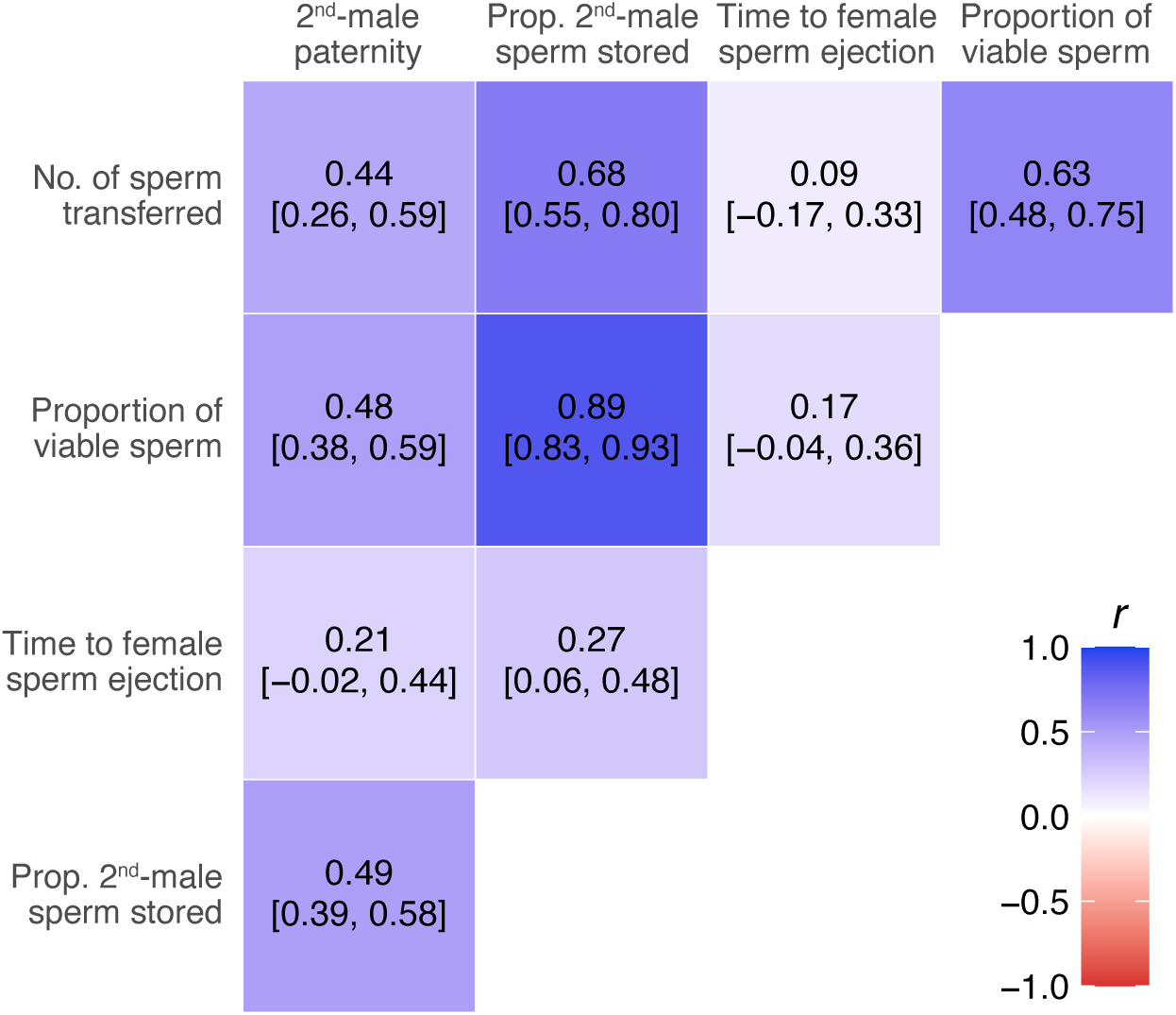
Correlations between the different phenotypic traits. Values reflect the averaged correlation coefficients with 95% confidence limits of pairwise Pearson correlations across bootstrapped group means (*N* = 1,000 samples) across males with varying microbiome consortia. Proportional traits (sperm viability, proportion of all stored sperm that came from the 2^nd^ male, and 2^nd^-male paternity shares) were logit-transformed before bootstrapping. Colors indicate the strength and direction of correlations as reflected by the color-gradient bar.

Moreover, higher-order interactions played a significant role in the expression of all traits, with significant five-way microbial interactions explaining variation in sperm viability (GLMM: *N* = 320, *OR* = 0.29 [0.13, 0.63]), the timing of female sperm ejection (LMM: *N* = 582, *β* = 150.10 [41.74, 255.22]), and the proportion of second-male sperm stored by females thereafter (GLMM: *N* = 582, *OR* = 2.34 [1.38, 4.11]) (Supplementary Figs. S9–S12). Despite the absence of a five-way interaction, the number of sperm transferred was explained by a four-way interaction of *Ao × Ap × Lb × Lp* (LMM: *N* = 582, *β* = 393.09 [198.49, 582.51]), alongside widespread non-additive effects at lower-order combinations (Supplementary Fig. S12).

Predictive models assuming additivity based on mono- or di-associations consistently underestimated observed values for sperm viability and storage, whereas deviations from additivity were weaker, though still detectable in several consortia, for sperm transfer and the time to female sperm ejection (Supplementary Fig. S13). Similarly, we also detected widespread “bystander” effects, in which the impact of a given microbial pair or triplet on traits was significantly modulated by the presence of additional species (Supplementary Fig. S14).

These results demonstrate that microbiome-mediated reproductive benefits emerge from nonlinear, composition-dependent effects on multiple ejaculate traits, each of which contributes to enhanced fertilization success.

## Discussion

Our findings revealed that the microbiome constitutes an overlooked but powerful determinant of fitness outcomes under sexual selection. Gut microbial composition, particularly the relative abundance of *Acetobacter* and species richness, predicted key reproductive traits and competitive fertilization success in *Drosophila melanogaster*. By manipulating microbiome composition across 32 unique consortia, we uncovered complex, non-additive, and context-dependent microbial effects that varied across reproductive traits. At the same time, broader patterns emerged: species richness strongly enhanced sperm viability and female sperm storage, whereas *Acetobacter* dominance specifically improved sperm transfer and second-male paternity success (Fig. 2). These trait-specific effects, which were robust across competitive contexts (axenic vs. conventional rivals) and multiple sensitivity analyses, point to distinct physiological mechanisms through which the microbiome influences post-mating outcomes.

### Proposed mechanistic pathways linking microbiome and reproductive traits

Microbial diversity likely enhances sperm viability and storage by mitigating oxidative stress and supporting sperm metabolic vigor. For example, *Drosophila* testes are characterized by high levels of the critical antioxidant glutathione, and of phosphoarginine, carnitine, and acetylcarnitine that are all essential for energy metabolism and sperm motility (Chintapalli *et al*., 2013; Zhang *et al*., 2020; Scolari *et al*., 2021). Testes also contain ether lipids (e.g., plasmalogens) (Chintapalli *et al*., 2013), which are essential for sperm membrane integrity (Reisse *et al*., 2001), and diverse gut microbiota may stabilize the redox balance or provision nutrients to support these metabolites and thus enhance the viability of sperm and their ability to displace resident sperm for abundant storage (Lüpold *et al*., 2012, 2020).

The positive effects of *Acetobacter* prevalence on sperm transfer and competitive paternity success may reflect acetate-mediated stimulation of insulin/insulin-like signaling (ISS) and host energy metabolism, both of which are known to be impaired in axenic flies (Shin *et al*., 2011; Kamareddine *et al*., 2018). This effect is likely amplified by larger ejaculates in *Acetobacter*-dominant males (e.g., by 4–28% relative to axenic males; Supplementary Fig. S7). While we used sperm quantity as a proxy, a larger ejaculate is expected to also contain a greater volume or altered composition of seminal fluid (Macartney *et al*., 2021; Zeender *et al*., 2023), which could directly influence fertilization capacity or female sperm use (Avila *et al*., 2011). Male accessory glands in *D. melanogaster* show high levels of methionine derivatives, most notably S-adenosylmethionine and its decarboxylated form, S-adenosylmethioninamine (Chintapalli *et al*., 2013). These derivatives serve as precursors to polyamines such as spermidine and spermine, which in turn act as critical regulators of sperm and seminal fluid function in diverse taxa (Lefèvre *et al*., 2011; Méndez, 2018; Scolari *et al*., 2021). We therefore propose that *Acetobacter*, through its acetate output, modulates host energy and amino acid metabolism and so indirectly enhances the capacity for polyamine synthesis in reproductive tissues. The direct production of polyamines by *Acetobacter* remains unconfirmed but is likely less relevant than this host-mediated effect.

Conversely, *Lactobacillus*-dominant consortia showed lower fitness gains relative to the axenic state (–3 to 21% for sperm transfer, 15–58% for sperm viability; Supplementary Fig. S7) and reduced focal-male paternity shares relative to *Acetobacter*-dominant consortia regardless of microbiome presence in the rival (0.68–0.74 vs. 0.78–0.87), which may be due to elevated reactive oxygen species (ROS) via DUOX activation that disrupt metabolic homeostasis (Ha *et al*., 2009; Iatsenko *et al*., 2018). Overall, our results align with conserved microbial roles across taxa (McFall-Ngai *et al*., 2013; Rowe *et al*., 2020; Li *et al*., 2025; Ramond *et al*., 2025) and generate specific, testable hypotheses: (i) that *Acetobacter*-derived acetate fuels polyamine synthesis in accessory glands to enhance competitive ejaculates, and (ii) that microbial diversity supports glutathione-dependent antioxidant defense in testes to ensure sperm integrity. These hypotheses can be tested by tracing isotopic labels from microbial metabolites to reproductive tissues and by assessing reproductive performance in flies with genetic disruptions in the key pathways of polyamine (e.g., *SamDC*) or glutathione synthesis (e.g., *Gclc*), or of IIS (e.g., *chico*).

### Microbiome-driven phenotypic variation and sexual selection

Our results broaden sexual selection theory by identifying the gut microbiome as a dynamic source of phenotypic variation in male reproductive fitness. This variation was primarily driven by the relative abundance of *Acetobacter*, which robustly predicted male competitive fertilization success (P_2_), rather than by the total number of bacterial species or their abundances. Unlike many other environmental factors, the microbiome functions as a dynamic, host-associated system that is reciprocally modulated by host ecology, behavior, and genotype while in turn amplifying or constraining host traits under selection (Spor *et al*., 2011; Koskella *et al*., 2017; Rowe *et al*., 2020). For instance, the 20–27% gains in paternity for diverse or *Acetobacter*-dominated microbiomes relative to germ-free males (Fig. 1B) suggest that beneficial consortia enhance competitive outcomes, providing an ecologically responsive axis of variation that may help maintain heritable differences in fitness-related traits despite strong directional selection (Rowe & Houle, 1996; Tomkins *et al*., 2004).

Given the prevalence of sperm competition across animals (Simmons & Fitzpatrick, 2012; Fitzpatrick & Lüpold, 2014) and the accumulating evidence for interactions between the gut microbiome and male reproduction in diverse taxa (McNamara *et al*., 2024; Zou *et al*., 2024; Ferenczi *et al*., 2025; Wu *et al*., 2025), our findings reinforce the need to evaluate the generality of microbially mediated effects on sexually selected traits. Comparative work could therefore test whether similar microbial consortia or metabolites influence competitive fertilization success and its associated traits across taxa, from model systems in basic research to applied breeding contexts. Such comparisons would move the field beyond documenting species-specific patterns toward identifying the principles that govern microbially mediated reproduction across animals.

### Evolutionary implications and future directions

Non-additive, context-dependent microbial interactions (e.g., pairwise effects modulated by bystander species; Fig. 3) indicate that microbiome complexity itself could be a target of selection. Host genotypes may evolve mechanisms, such as immune modulation or behavioral transmission, to favor high-performing communities like *Acetobacter*-dominated ones (see above) (Foster *et al*., 2017; Moran *et al*., 2019). This establishes the microbiome as an extended reproductive phenotype with potential feedback into host evolution (Koskella & Bergelson, 2020; O’Brien *et al*., 2024). Furthermore, female responses, such as delayed sperm ejection and biased storage of microbiome-enhanced ejaculates, suggest at least partially microbiome-mediated cryptic female choice (Rowe *et al*., 2020), where microbial signals in seminal fluid could influence post-copulatory processes (Eberhard, 1996; Firman *et al*., 2017; Daupagne *et al*., 2025). While this remains to be experimentally tested, it aligns with microbiome effects on female reproduction in other taxa (Chadchan *et al*., 2022) and warrants exploration in natural populations. To address potential limitations, future work should explore diet–microbiome synergies or interactions with infections, which could further modulate these effects.

Recognizing that microbial partners can dynamically alter reproductive trait expression and competitive fertilization success, our results provide empirical support for the view that host–microbe interactions represent a third axis of sexual selection, alongside genetics and the environment (O’Brien *et al*., 2024). This perspective broadens sexual selection theory and motivates four key research priorities: (i) mechanistic studies using metabolomics and transcriptomics in gnotobiotic models to quantify microbial outputs (e.g., acetate) and their effects on the function of host reproductive tissues, thereby extending gut-focused studies (Douglas, 2018; Matthews *et al*., 2021); (ii) characterizing transmission modes—vertical, social, or sexual—to clarify how beneficial consortia persist and contribute to heritable variation (Koskella *et al*., 2017; Leftwich *et al*., 2017); (iii) testing male–female microbiome interactions in non-axenic females to assess compatibility effects on sperm handling or processes of cryptic female choice (Rowe *et al*., 2020); and (iv) ecological and comparative studies of diet-microbiome synergies, genotype-microbiome epistasis, and cross-taxa patterns with conserved guilds (e.g., acetate producers) to establish generality (Broderick & Lemaitre, 2012; O’Brien *et al*., 2024; Araujo *et al*., 2025).

### Conclusion

This study establishes the gut microbiome as a causal and evolutionarily relevant factor underlying competitive fertilization success in *D. melanogaster*, driven by emergent, non-additive interactions within microbial communities. By demonstrating that *Acetobacter* dominance and species richness shape distinct ejaculate traits and female sperm use, we bridge microbial ecology with core concepts in sexual selection. Our findings reveal the microbiome as a critical source of phenotypic variation and expand our understanding of reproductive trait evolution (Rosenberg & Zilber-Rosenberg, 2018; O’Brien *et al*., 2024). This integration of host–microbe partnerships into sexual selection theory opens new avenues for exploring coevolution, the maintenance of heritable trait variation under strong selection, and the mechanistic pathways involved, while also motivating parallel work in natural populations to determine how these interactions shape reproductive outcomes in the wild.

## Materials and Methods

### Experimental populations

Throughout this study, we used outbred populations of *Drosophila melanogaster* (LH_m_ strain). These were either derived from an outbred wild-type population or, for standard competitors in the sperm competition assay, from a backcrossed population that had been genetically modified to ubiquitously express green fluorescent protein (GFP), permitting paternity assignment against the focal wild-type males (Lüpold *et al*., 2012, 2013). To study biases in female sperm storage, we used focal males whose sperm heads expressed red fluorescent protein (RFP), and our standard competitors with ubiquitous GFP expression, permitting quantification of each competitor’s sperm inside the female reproductive tract (Manier *et al*., 2010; Lüpold *et al*., 2012, 2013). We maintained all fly populations at moderate densities under constant conditions of 25°C, 60% humidity, and a 12h:12h light:dark cycle.

### Generating axenic flies

To generate axenic flies, we performed all steps hereafter in a laminar flow chamber to avoid contamination. First, we collected eggs from Petri dishes containing grape-juice agar with a smear of yeast paste that had been placed in four large population cages after a six-hour egg-laying period. After washing these eggs off the plates into a strainer, we dechorionated them with 3.5% bleach for 2 minutes, followed by a rinse in 70% ethanol and three rinses with sterile water (Koyle *et al*., 2016). We then transferred these eggs to autoclaved standard fly food supplemented with antibiotics (ampicillin, kanamycin, erythromycin, and tetracycline at final concentrations of 50, 50, 10, and 10 μg/mL of food medium, respectively) in autoclaved glass vials. These vials were then left undisturbed under standard conditions (100 embryos/vial, 25°C, 60% humidity, and a 12h:12h light:dark cycle) throughout development.

Upon fly eclosion, we collected both males and females as unmated individuals, separated them by sex and confirmed their axenic state by plating a homogenized subsample of flies on LB and MRS agar followed by 16S PCR. Briefly, we extracted genomic DNA from 3−5 pooled axenic flies per sample using the ZymoBiomics^TM^ DNA Microprep kit following the instructions provided by the manufacturer. We also extracted DNA from flies with an intact microbiome to serve as positive controls. For DNA amplification, we used bacterial universal primers 27F and 1492R as forward and reverse primers, respectively. We prepared the PCR master mix using 9.5 μL of nuclease free water, 12.5 μL kappa HiFi master mix, 1 μL of forward and reverse primer and 1 μL of template DNA. The PCR cycle included an initial denaturation at 95°C for 90 s, followed by 35 cycles of denaturation at 95°C for 30 s, annealing at 55°C for 30 s, an extension at 72°C for 90 s, and a final extension at 72°C for 5 mins. We ran the PCR products on a 1% agarose gel and visualized them under ultraviolet light after staining with SYBR Safe (Invitrogen) to confirm the absence of bacteria in axenic individuals and the presence of bacteria in controls.

We maintained all females and standard competitors as axenic flies throughout the experiment and supplemented axenic focal males with different bacterial cultures to produce gnotobiotic associations as described below. All adult food was free of antibiotics.

### Bacterial cultures and gnotobiotic associations

Throughout this study, we used five bacterial species isolated from the gut of wild-type flies (Gould *et al*., 2018): *Acetobacter orientalis*, *A. tropicalis*, *A. pasteurianus*, *Lactobacillus plantarum*, and *L. brevis*. We associated our focal flies with all 32 possible combinations among these bacterial strains following the general protocol of Gould et al. (2018). To prepare bacterial cultures, we grew them overnight: *Lactobacillus* spp. in MRS (De Man–Rogosa–Sharpe) broth without shaking and *Acetobacter* spp. in MRS + mannitol medium at 30°C on a shaker. The next morning, we checked these bacterial cultures for growth and performed their optical density (OD) measurements at 600 nm. Then, we spun down these different bacterial cultures at 7000 rpm for 5 minutes and resuspended the resulting pellets to a concentration of 10^8^ cells/mL in sterile phosphate-buffered saline (PBS). Resuspension volume was determined following established protocols (Newell & Douglas, 2014; Koyle *et al*., 2016). Finally, we added a 50 µL (i.e., ca. 5×10^6^ colony forming units, CFUs) of bacterial culture to sterile (but antibiotic-free) fly food vials, adjusting the volume per bacterial strain in multi-species associations for equal representation within a consistent total density, which itself was comparable to prior reports for conventional flies (Newell & Douglas, 2014; Gould *et al*., 2018).

After adding the bacteria to food vials and air-drying these under in laminar flow hood, we transferred 10 one-day-old axenic and unmated males to each vial for inoculation with the added bacterial strains. We moved these males to fresh food vials with same bacterial treatment every alternate day for 5 days. Thereafter, we confirmed gnotobiotic associations by plating together the homogenates of 3-4 adult flies from same treatments and checking their colony morphologies, and by Sanger sequencing of 16S PCR homogenates. For multi-associations, we plated the fly homogenates and picked the distinct types of colonies from a given plate for PCR and performed sequencing to confirm that morphologically similar species corresponds to same species in all the replicates, thus allowing us to distinguish between different bacterial species by looking at their colony appearance. Thereafter, we tested the presence of the expected bacterial species in a particular bacterial combination by plating them and looking at the colony morphologies.

### Bacterial abundance in different combinations

To determine bacterial abundance in different gnotobiotic associations, we counted the CFUs per fly for each bacterial species across the different gnotobiotic associations. To this end, we washed males with 70% ethanol, individually homogenized them with a pestle in microcentrifuge tubes, and then diluted these homogenates to different dilutions with a final volume of 100 μL to find a dilution with comparable colony counts (approx. 100 CFUs) for visual quantification. In gnotobiotic cultures with both bacterial genera, we transferred an aliquot of well-mixed fly homogenate to two plates, supplementing the MRS media of one plate with 10 μg/mL ampicillin to selectively inhibit the growth of *Lactobacillus*, while incubating the other in a CO_2_ atmosphere to prevent the growth of *Acetobacter*. We achieved within-genus differentiation of strains by colony morphologies: *A. orientalis* with flat colonies with ruffled borders, *A. tropicalis* with small tan colored and highly reflective colonies, and *A. pasteurianus* with large tan colonies. Similarly, *L. brevis* produces brownish yellow colonies and *L. plantarum* milky white colonies (Supplementary Fig. S15). Using 16S PCR amplification followed by Sanger sequencing, we confirmed specificity of these colony properties, demonstrating the robustness of our method based on morphological identifications during experiments. After counting the number of colonies on each plate, we calculated CFU per fly as CFU = *C* × *D* × *V* / *P*, where *C* = number of colonies counted, *D* = dilution factor, *V* = volume of fly homogenate, and *P* = volume plated (Koyle *et al*., 2016).

Across all 10 replicate plates combination examined, the expected species were present, and none of the remaining species. Consistent with Gould et al. (2018), total CFU counts increased with species richness while the species-specific abundances declined (Supplementary Fig. S2).

### Sperm competitive ability

To investigate sperm competitiveness of focal males, we quantified their sperm offense ability (P_2_) by mating them as the second males with axenic females that, three days earlier, each had been inseminated by a GFP-labeled male with no (axenic) or a conventional (unmanipulated) microbiome. We monitored the occurrence of copulations for 7-8 hours and transferred remated females to oviposition vials while giving those that did not remate another two 8-hour remating opportunities on the following days. Flies that did not remate by the last opportunity (within these 3 days) were discarded. We allowed all remated females to lay eggs for three days. After all progeny eclosed, we scored them for paternity under a fluorescence stereoscope, based on the presence/absence of the ubiquitous GFP marker in standard competitors. The proportion of second-male progeny indicated the focal-male sperm competitive ability (i.e., sperm offense). Against both axenic and conventional competitors, we paired 30 experimental males per consortium across three blocks, resulting in a total of *N* = 960 trials in each experiment.

### Sperm viability

We quantified sperm viability for 10 males per bacterial treatment (total *N* = 320 males). Once males were six days old and inoculated, we anaesthetized them on ice and dissected them on a microscopic slide to isolate the seminal vesicles from their reproductive tissue. We then placed one of the seminal vesicles in 10 µL of Ringers solution and punctured it with an insect pin to release the sperm mass for immediate staining with LIVE/DEAD fluorescent dye, following the principles of prior studies on *Drosophila* (Tourmente *et al*., 2019; Meena *et al*., 2024). This two-component dye consists of SYBR-14 (‘LIVE’ component emitting green fluorescence), which is actively incorporated into living cells, and propidium iodide (PI, with red fluorescence) that can only permeate damaged membranes (typical of dead cells). We prepared an aqueous staining solution by combining 2.4 µL of SYBR-14 stock solution (1 mM in DMSO) and 4.5 µL of PI stock solution (2.4 mM in water) in 93.1 µL of Ringer’s solution. Then, we added 5 µL of the staining solution to the sperm sample before covering it with a coverslip and incubating it for 5 minutes in the dark. Under an Olympus BX50 fluorescence microscope with a GFP/RFP dual filter, we captured images of randomly chosen areas of each sample to later manually count at least 200 sperm and calculate the proportion of viable sperm in ImageJ v. 1.34 (Schneider *et al*., 2012).

### Visualizing sperm competition within female reproductive tract

Since competitive fertilization events ultimately are the result of both male and female effects (Eberhard, 1996; Lüpold *et al*., 2020), we investigated how females bias their sperm storage based on interactions with males supplemented with different microbiome. *For this experiment, we again paired* axenic unmated females first with an axenic GFP male and, three days later, with a focal male as for sperm competitive ability (above). To visualize sperm of both males, we used focal males expressing RFP in their sperm heads. Immediately after remating, we transferred females to individual wells of cell-culture plates, with a coverslip held in place with dots of rubber cement in two corners. We checked these females every 10 min for up to 6 h for the ejection of the sperm mass containing both displaced first-male and excess second-male sperm (Manier *et al*., 2010; Lüpold *et al*., 2012). The timing and extent of sperm ejection have previously been reported to affect relative numbers of first- and second-male sperm stored by females and, consequently, competitive fertilization success among males (Lüpold *et al*., 2013, 2020).

Upon female sperm ejection, we recorded the interval from mating to ejection before removing females from the well and freezing them individually in Eppendorf tubes. Thereafter, we dissected these females and counted the green (standard competitors) and red sperm (focal males) within the female reproductive tract (paired spermathecae, seminal receptacle and bursa copulatrix) under a fluorescence microscope. From these counts, we determined S_2_, the proportion of all sperm stored by the female that came from the second (experimental) male. We also counted the respective sperm numbers in the ejected sperm mass to estimate, combined with above counts, the total number of sperm transferred by the second male and the number of first-male sperm still residing in storage at the time of remating. We performed this experiment in five blocks with 10 replicates per treatment and block (total *N* = 1,600 remating trials).

### Statistical analyses

We conducted all analyses in R v4.4.3 (R Core Team 2025), using bootstrapping procedures with 1,000 iterations to assess the stability and robustness of model estimates. To evaluate the effects of microbial diversity on reproductive outcomes, we resampled microbiome compositions (*N* = 32 unique consortia) and fitted mixed-effects models across all bootstrap datasets. These models included species richness (number of bacterial species) as a fixed effect and both experimental block and microbial composition (i.e., consortium identity) as random effects. Depending on the response variable, models were either linear mixed-effects models (for traits such as the number of sperm transferred and time to sperm ejection) or generalized linear mixed-effects models with a binomial error distribution and logit link function (for proportions of viable sperm, second-male sperm in storage, and second-male progeny [P_2_]). Binomial models incorporated an observation-level random effect to account for overdispersion.

To further evaluate the combined effects of species richness, total bacterial load, and the proportional representation of *Acetobacter* in each consortium (both latter variables based on CFU counts extracted from homogenized males), we fitted non-mixed models across bootstrapped datasets, as bacterial counts and phenotypic traits were assessed in separate fly cohorts under identical conditions due to logistical constraints. These datasets were independently bootstrapped and combined for analysis. To address the collinearity between species richness and bacterial load, we conducted a series of sensitivity analyses within our bootstrapped framework (detailed in the *Supplementary Material*), which confirmed the robustness of our estimates.

To examine the contribution of specific microbial interactions, we fitted mixed-effects models using the presence/absence of each bacterial species and their two- to five-way interactions as fixed effects, with block and, where appropriate, observation identity as random effects. In this interaction-focused analysis, we bootstrapped at the level of individual observations, stratified by microbial consortium, to retain the structure of specific combinations. All subsequent analyses used the same set of bootstrapped datasets for consistency and comparability.

To better understand the importance of species combinations in the microbiome and to isolate emergent effects beyond individual species contributions, we used an epistasis-based framework to elucidate how well the averaged phenotypic effects of single- or pairwise species associations predict higher-order phenotypes (Gould *et al*., 2018). For single-species predictions, we compared the measured trait values of males associated with two- to five-species microbiotas to values averaged across mono-associated males that jointly represented the same bacterial combinations. We then plotted these differences (averaged prediction minus measured trait value) with bootstrapped 95% confidence intervals to visually determine if predictions deviated from the measured estimates (i.e., confidence interval excluding zero difference), and repeated this procedure to predict phenotypes by averaging the estimates of di-associated males (for calculations see Gould *et al*., 2018).

Following the same general principle, we further quantified how the presence or absence of non-focal (“bystander”) community members modulates both pairwise and three-way microbial interactions by exhaustively enumerating all possible bystander contexts. We compared this “contextual” analysis against a “standard” test to reveal how higher-order community background shapes emergent effects on host traits. Following Gould et al.’s (2018) nomenclature, the “standard” test calculated the interaction coefficient among all species present in a specific community configuration, whereas the “contextual” tests assessed the interaction between a focal species pair within the context of additional bystander species. We computed interaction coefficients for each focal species pair or triplet across all possible presence/absence combinations of bystander species and plotted them with bootstrapped standard errors to highlight the varying directions of interactions depending on the bystander combinations. In both the single/double-species predictions and these standard/contextual tests, we derived the coefficients to calculate the deviations from a mixed-effects models with an appropriate link function for each microbial combination, controlling for block effects. We repeated each model across 1,000 bootstrapped datasets to calculate the standard errors or 95% confidence intervals from the distribution of the calculated coefficients or interaction differentials.

## Acknowledgments

We thank W. Ludington for providing the five bacterial strains used in our experiments, and the Swiss National Science Foundation (grant PP00P3_202662 to S.L.) for financial support.

## Author Contributions

K.M.: conceptualization, methodology, investigation, data collection and curation, data analysis, writing original draft, writing—review and editing, project administration; A.M.: investigation, data collection, writing—review and editing; A.N.D.N: investigation, data collection, writing—review and editing; S.H.S: investigation, writing—review and editing; J.R.: investigation; S.L.: conceptualization, methodology, funding acquisition, data analysis, project administration, supervision, writing original draft, writing manuscript—review and editing. All authors gave final approval for publication and have agreed to be held accountable for the work performed therein.

