## Supplemental Methods, Figures and Tables for "Male gut microbiome mediates post-mating sexual selection in *Drosophila*"

### Supporting Text

#### Sensitivity analyses

**General description.** To ensure the robustness of our findings linking gut microbiome composition (particularly bacterial species richness, total bacterial load, and the relative abundance of *Acetobacter*) to male competitive fertilization success ($P_2$) and associated host ejaculate traits, we conducted a comprehensive suite of sensitivity analyses. Given that no single method for addressing multicollinearity is universally superior and each has inherent limitations (e.g., potential bias in residualization, shrinkage in regularization, or sensitivity to assumptions in simulations (Dormann *et al.*, 2013)), we employed a complementary multi-method approach to mitigate these issues and strengthen confidence in our conclusions. These analyses specifically targeted potential multicollinearity among predictors (e.g., between species richness and bacterial load; Pearson's $r \approx 0.45$–$0.60$ across bootstraps), which could inflate variance estimates or bias coefficients in multivariate models. Multicollinearity arises when predictors are correlated, leading to unstable estimates, inflated standard errors, and difficulty in interpreting individual effects (Graham, 2003; Dormann *et al.*, 2013).

Our approach combined orthogonalization techniques (residualization), data subsetting, variable exclusion, regularization (elastic net including ridge), variance partitioning (commonality analysis), and simulation-based validation. We performed all analyses within a bootstrapped framework ($N$ = 1,000 resamples with replacement) to account for sampling variability and provide distribution-free confidence intervals (Efron & Tibshirani, 1994). We fitted models using generalized linear models (GLMs) for binomial responses (i.e., proportional sperm representation in female storage, sperm viability, and paternity shares) or linear models (LMs) for continuous traits (e.g., sperm transfer or time to female sperm ejection), with predictors standardized within bootstraps to facilitate comparability (mean = 0, SD = 1). For GLMs, we used a logit link and checked for overdispersion; for binomial traits with weights (e.g., total progeny for $P_2$), we incorporated weights to reflect sample size per observation.

We implemented all models in R v. 4.4.3, with key packages including *dplyr* (Wickham *et al.*, 2025) for data manipulation, *broom* (Robinson *et al.*, 2025) for tidying outputs, *car* (Fox & Weisberg, 2019) for VIFs, *DescTools* (Signorell, 2025) for pseudo-$R^2$, *glmnet* (Friedman *et al.*, 2010) for elastic net/ridge, *MASS* (Venables & Ripley, 2002) for multivariate normals, *ggplot2* (Wickham, 2016), *ggridges* (Wilke, 2025), and *patchwork* (Pederson, 2025) for visualizations (e.g., ridge plots, bar plots with CIs, density comparisons).

These analyses collectively validate our core conclusions that: (i) microbiome diversity (species richness) and *Acetobacter* dominance positively influence $P_2$ and its underlying reproductive traits, (ii) effects are non-additive and community-driven, and (iii) bacterial load has secondary, context-dependent roles. The broad consistency across all methods strongly indicates that the results are not mere artifacts due to collinearity.

**1. Residualization to orthogonalize collinear predictors.** Residualization removes shared variance between collinear predictors, thereby creating orthogonal terms that isolate unique effects without discarding data (Freckleton, 2002; Dormann *et al.*, 2013). This method is particularly useful in our study where bacterial load and richness were inherently linked (e.g., due to microbial growth dynamics), though it can introduce slight bias if the auxiliary regression is mis-specified or if the residuals do not fully capture nonlinearity (Freckleton, 2002; Warton *et al.*, 2015). We residualized richness on load (or vice versa) to test if effects persist after accounting for their covariance. For each bootstrap, we computed residuals from linear regressions of standardized richness on load or load on richness. These residuals replaced the original terms in multivariable models, alongside the proportional representation of *Acetobacter*. We fitted LMs for continuous or binomial GLMs (logit link) for proportional responses, respectively. Using *broom::tidy()*, we then extracted effect sizes: correlation coefficients ($r = t/\sqrt{t^2 + df}$) for LMs and odds ratios ($OR = e^{\beta}$) for GLMs. Across all responses, the effect sizes remained largely stable relative to the original models. All VIFs dropped to ≤1.18 (Tables S1 and S2), well below the recommended threshold of 5 (O'Brien, 2007), thereby confirming the reduction of collinearity. These results aligned with the original models and affirm that the microbiome effects are not mere artifacts of collinearity but reflect distinct biological pathways (Douglas, 2015).

**2. Subsetting to restrict predictor range.** Collinearity often amplifies in datasets with extreme values or outliers. Subsetting to a narrower range (e.g., within 1 SD of the mean) can reduce variance inflation while preserving central tendencies, though it reduces sample size and power (Belsley *et al.*, 1980; Dormann *et al.*, 2013). This conservative approach tests effect robustness in a "low-collinearity" subsample, focusing on typical microbiome compositions. Within each bootstrap, we subset data to samples where standardized bacterial load fell within ±1 SD of its mean (retaining ~74% of data, i.e. = 23 of the 31 microbiome compositions per bootstrap). The models then excluded load but retained both richness and *Acetobacter* proportion on the reduced datasets. The effects largely mirrored the original models (Table S3). Coverage rates (proportion of bootstraps with valid fits) were 100%, and the subset sample size was sufficient (>15 per fit). Hence, the main effects were not driven by outliers, thereby validating our results that microbiome community composition modulates post-mating success.

**3. Variable exclusion to isolate effects.** Excluding one collinear predictor per model isolates the influence of others, providing a simple diagnostic for confounding (Graham, 2003). This method is akin to sensitivity testing in meta-analyses (Viechtbauer & Cheung, 2010) and helps discern if effects are load- vs. richness-dominated, though it risks omitting relevant variables and underestimating shared variance. We fitted models as in section 1, but this time including only species richness *or* bacterial load alongside the proportion of *Acetobacter*. Exclusion did not alter key patterns, in that the effects of the relative abundance of *Acetobacter* and species richness remained qualitatively unchanged when bacterial load was excluded (Table S4), and the same was true for the remaining variables when excluding species richness (Table S5).

**4. Ridge regression for regularization.** Ridge regression adds a penalty (λ) to shrink coefficients, thereby stabilizing estimates under multicollinearity without bias in large samples (Hoerl & Kennard, 1970; Dormann *et al.*, 2013). This can help where predictors may share metabolic pathways (e.g., acetate production by *Acetobacter* influencing load/diversity (Chaston *et al.*, 2014)), though it introduces bias toward zero and requires λ tuning. A limitation is that ridge does not perform variable selection (unlike lasso, α = 1), so it retains all predictors but reduces their influence if collinear.

Using the *glmnet* package (Friedman *et al.*, 2010), we applied elastic net penalties, with α = 0 for ridge, α = 0.2 for mixed, and α = 1 for lasso, to standardized predictors in regularized regression frameworks within each bootstrap replicate. For proportional responses (e.g., $P_2$), we logit-transformed *y* to handle bounded data while modeling as gaussian. This quasi-likelihood approximation accommodated the non-integer values arising from bootstrap averaging (where resampling with replacement and aggregation per treatment group yielded fractional "counts" that were unsuitable for strict integer-based binomial modeling). To integrate with the existing code structure (section 1), we converted the standardized predictors to a matrix and selected λ via 5-fold cross-validation in *cv.glmnet* to minimize mean squared error or deviance, using *lambda.min* as the penalty parameter. We skipped occurrences of zero variance in *y* or predictors and then derived confidence intervals from bootstrap distributions of the coefficients for each α value. Penalization minimally altered estimates, with minimal shrinkage for dominant predictors like the proportional representation of *Acetobacter* (Table S6), thus confirming stability of the key microbiome effects.

**5. Variance partitioning.** To disentangle the unique and shared contributions of *Acetobacter* abundance, species richness, and total bacterial load, we applied variance partitioning (a form of commonality analysis (Ray-Mukherjee *et al.*, 2014)). In host-microbe systems, this can disentangle community-level synergies (e.g., richness amplifying *Acetobacter* benefits (Wong *et al.*, 2013)). Commonality analysis decomposes explained variance into unique and shared fractions, thereby revealing additivity or interactions, though it can be sensitive to collinearity and model misspecification (e.g., suppression effects leading to negatives) (Graham, 2003; Ray-Mukherjee *et al.*, 2014).

For each bootstrap replicate, we fit models to all subsets of predictors and used inclusion–exclusion formulas to calculate the variance uniquely attributable to each predictor, as well as variance shared among predictors (Peres-Neto *et al.*, 2006; Legendre, 2008). For linear models, we used adjusted $R^2$ as the measure of explained variance, and for logistic models we used McFadden's pseudo-$R^2$ (McFadden, 1974). Because commonality analysis can yield negative shared fractions when predictors are highly collinear, we report two complementary versions: (i) the true decomposition, which includes negative fractions and directly reflects redundancy among predictors, and (ii) a cleaned version, in which negative fractions were truncated at zero and values rescaled to sum to 100% of the explained variance. Both approaches gave qualitatively consistent conclusions, but the cleaned version aids visualization and interpretability. Overall, we decomposed the total variance into empty ($R^2_0$), single ($R^2_{A/B/C}$), pairwise ($R^2_{AB/AC/BC}$), and full ($R^2_{ABC}$) partitions, which then allowed us to calculate unique ($U_A = R^2_A - R^2_0$, etc.) and shared ($S_{AB} = R^2_{AB} - U_A - U_B - R^2_0$, etc.) partitions, with percentages if $R^2_{ABC} > 0$.

Unique *Acetobacter* contributions dominated (e.g., $P_2$: 82–93%; sperm viability: 4–7%), with richness adding 8–17% uniquely for $P_2$ (Tables S7 and S8). Shared variances were low (<5% for most, except the time to sperm ejection where overlaps reached 14–42%), indicating minimal confounding. Negative shared terms in "true" partitions reflected suppression effects but did not undermine positives. This decomposition validates non-additive interactions, as unique components align with the original effects and support microbiome ecology as a driver of ejaculate variation (Borcard *et al.*, 1992; Legendre, 2008).

**6. Simulation-based validation of estimator performance.** We used simulation-based validation to evaluate the performance of our regression estimators—specifically their bias, mean squared error (MSE), and confidence interval (CI) coverage—under conditions that closely mimicked the empirical collinearity among the microbiome predictors (Morris *et al.*, 2019). This forward validation tested if the fitted models could recover the true effect sizes when predictors were correlated (Kauermann & Carroll, 2001), as in the observed data, and whether inference remained valid under small-sample uncertainty and predictor collinearity.

For each reproductive trait, we simulated 1,000 independent datasets using the empirical means and standard deviations of the fitted slopes from the base models as the true generative parameters. We drew predictors from a multivariate normal distribution (*MASS::mvrnorm*) with mean zero, unit variance, and appropriate intercorrelations. To match empirical correlations, we sampled bacterial load and species richness with an inter-predictor correlation of $\rho = 0.51$, while generating the proportional abundance of *Acetobacter* independently. The sample size ($N = 31$) reflected the microbiome-treated groups in the empirical data (i.e., excluding axenic flies that lacked a defined *Acetobacter* proportion).

We analyzed each dataset using the same model family (Gaussian or binomial) and structure as in the main analysis. We employed three inference methods: (i) Wald CIs (standard asymptotic normal intervals), (ii) HC3 heteroskedasticity-consistent CIs, which adjust for small-sample bias and model misspecification, and (iii) parametric bootstrap CIs (200 resamples per simulation), which re-estimate confidence limits from resampled response variables conditional on fitted parameters. However, we focused on HC3 in the final analyses due to its superior and stable coverage properties across conditions.

For each coefficient ($\beta$), we calculated across all simulations the bias ($= \mathbb{E}[\beta_{\text{estimated}} - \beta_{\text{true}}]$), mean squared error ($= \mathbb{E}[\beta_{\text{estimated}} - \beta_{\text{true}}]^2 = \text{bias}^2 + \text{Var}(\beta)$), and CI coverage ($= P(\beta_{\text{true}} \in [CI_{\text{lower}}, CI_{\text{upper}}])$), where we approximated expectations ($\mathbb{E}[\ldots]$) by the empirical mean over 1,000 simulation replicates. Thus, bias represents the average deviation of the estimated coefficient from its true value, MSE reflects the combined contribution of estimator bias and variance (i.e., total estimation error), and CI coverage denotes the proportion of simulations in which the 95% CI contained $\beta_{\text{true}}$.

Across all simulation scenarios, estimator performance was robust (Fig. S3). Bias remained low in all but two cases (|bias| < 0.04 for most predictors, typically < 0.02), indicating that the fitted models recovered the true effect sizes ($\beta_{true}$) with minimal systematic deviation. Mean squared error (MSE) values ranged from approximately 0.06–0.16 for Gaussian and 0.34–0.52 for binomial outcomes, consistent with modest estimator variance relative to the scale of the true effects (Table S9). This pattern reflects higher estimator variance in logistic regressions, as expected given their discrete outcomes and nonlinear link, whereas linear models yielded more stable estimates around the true effects.

For most predictors, 95%CI coverage approached or exceeded the nominal level (≥0.95; Fig. S3), confirming that uncertainty was well-calibrated. The only notable exceptions were effects of species richness and bacterial load on sperm transfer and ejection time, respectively, which showed mild undercoverage (0.78–0.87). To further assess their robustness, we extended simulations under controlled departures from model assumptions by varying the strength of heteroskedasticity ($\gamma$ = 0–0.6) and the tail heaviness of the error distribution [Student's *t*-distribution with 3, 5, or 10 degrees of freedom: *t*(df = 3, 5, 10)], where increasing $\gamma$ inflates variance with predictor magnitude while lower *t* degrees of freedom generate heavier-tailed residuals with greater kurtosis. For both response variables, HC3 confidence intervals maintained nominal or near-nominal coverage (≥0.9) and negligible bias even under the strongest tested distortions [$\gamma$ = 0.6, *t*(df = 3)] (Fig. S8). Interestingly, slightly lower coverage occurred under moderate distortions ($\gamma$ = 0.2–0.3), likely because partial heteroskedasticity introduces uneven leverage that is not yet fully compensated by the HC3 small-sample correction. At higher $\gamma$, the robust variance correction more effectively compensates to restore near-nominal coverage.

Together, these results confirm that the fitted models were unbiased, precise, and inferentially reliable under realistic sampling, collinearity, and moderate model misspecification.

**Figures**

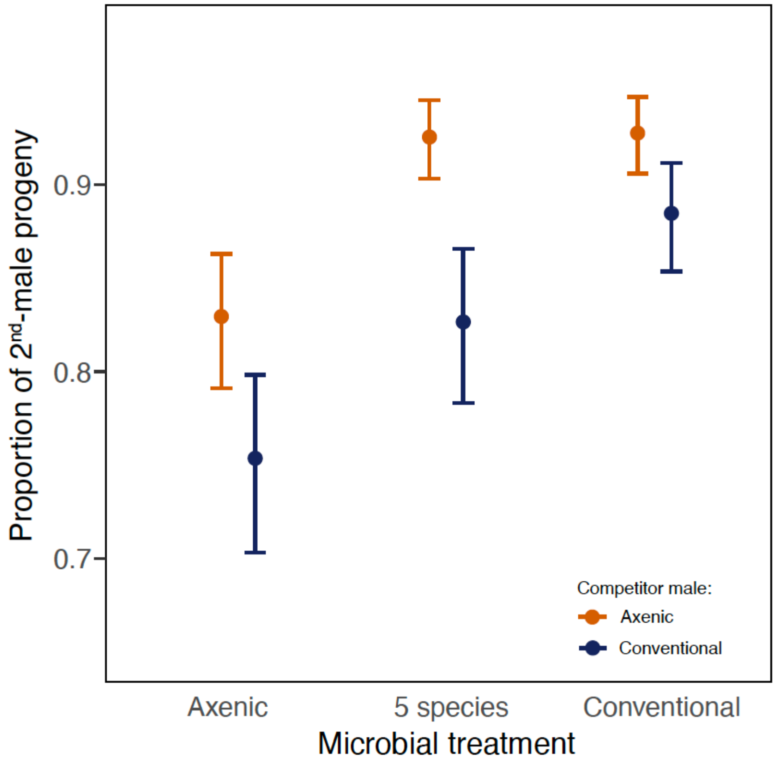


**Fig. S1.** Competitive fertilization success, measured as the proportion of second-male (i.e., focal-male) paternity, against either axenic or conventional competitors (first males to mate). In either context, males that were associated with the five-species microbial consortium (*Acetobacter orientalis*, *A. tropicalis*, *A. pasteurianus*, *Lactobacillus brevis* and *L. plantarum*) after axenic development outperformed consistently germ-free (axenic) males and even matched males with a conventional (entirely unmanipulated) microbiome in their sperm competitiveness, at least against germ-free competitors. Error bars represent 95% confidence intervals.

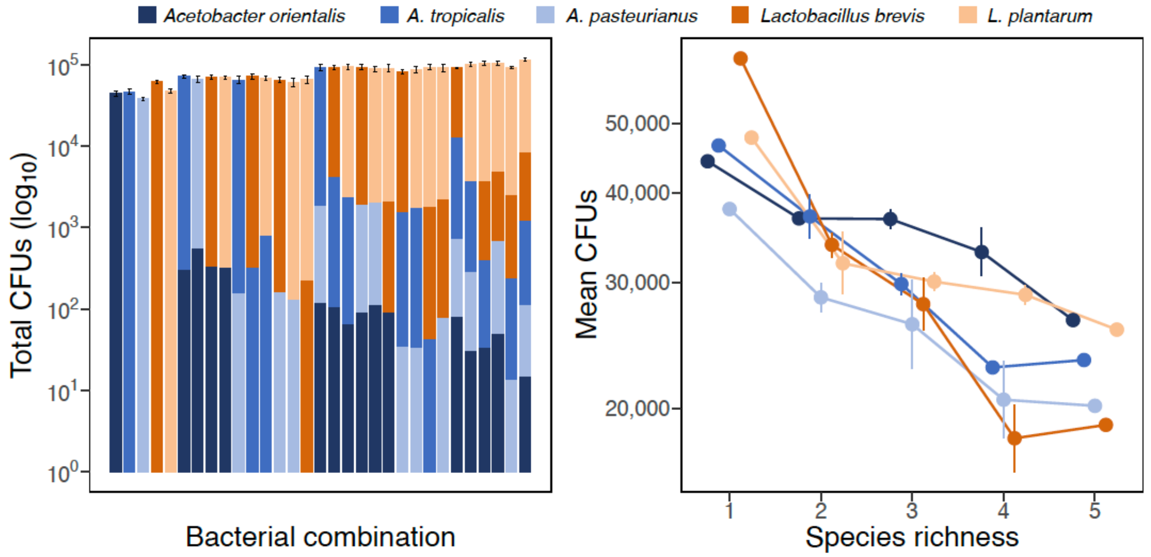


**Fig. S2.** Abundances of the different species (linear scale) for all 32 possible species combinations (*N* = 10 replicate flies per combination) with increasing species richness. The height of the bars in the left panel reflect the total microbial load ($log_{10}$ scale), with the relative abundances of the constituent species shown by coloration (linear scale). Total increase in total bacterial load with increasing species richness is shown by increasing bar height from left to right. The right panel depicts the decline in species-specific colony-forming units (CFUs) counts in response to increasing species richness. All error bars show 95% confidence intervals.

**Fig. S3 (next page). Simulation-based validation of model performance across reproductive traits.** Simulations ($n$ = 1,000 datasets per trait) assessed the accuracy and reliability of fitted models under realistic predictor collinearity ($\rho$ = 0.51 between bacterial load and species richness) and sample size ($N$ = 31). For each response variable, the left panels show the recovery of estimated slopes ($\beta$) for the proportion of *Acetobacter*, bacterial load, and species richness: colored density curves represent distributions of simulated estimates (light) and empirical bootstrap estimates (dark). The middle panels summarize estimator bias (mean [$\beta_{estimated} - \beta_{true}$]), and the right panels indicate the coverage rates of 95% confidence intervals, based on HC3 confidence intervals. Except for two cases (species richness in C and D), |bias| was <0.04 and generally < 0.02), and coverage rates exceeded 0.95 for all predictors in all simulations except for E and F, where it approximated nominal levels only for the proportion of *Acetobacter* (also see Table S9). Hence, these simulations confirmed that fitted models, in general, reliably recovered true effects even under empirical collinearity, thereby supporting the robustness of microbiome–fertility inferences reported in the main text.

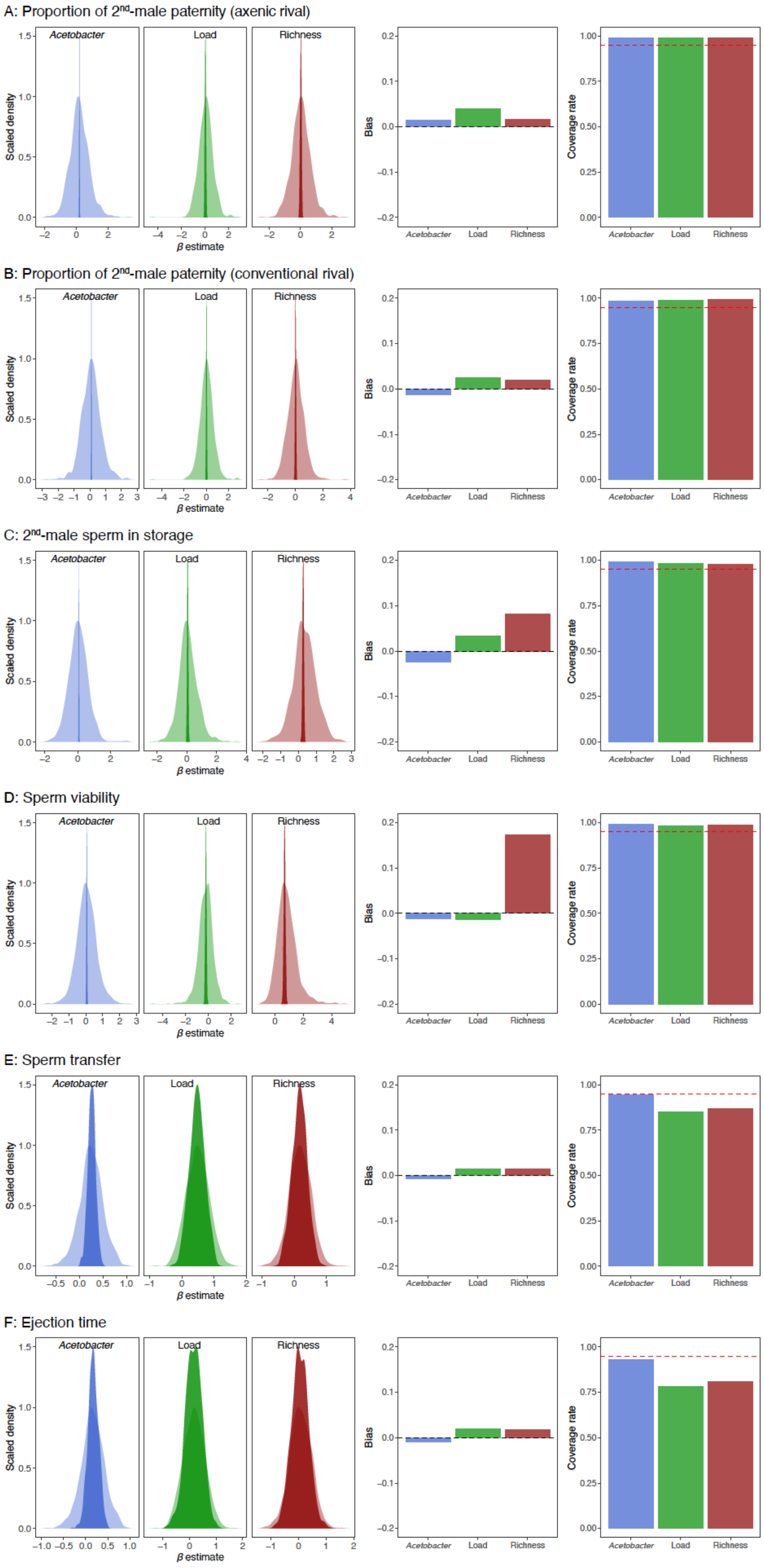
A: Proportion of 2nd-male paternity (axenic rival)
B: Proportion of 2nd-male paternity (conventional rival)
C: 2nd-male sperm in storage
D: Sperm viability
E: Sperm transfer
F: Ejection time
Acetobacter
Load
Richness
Scaled density
β estimate
Bias
Coverage rate

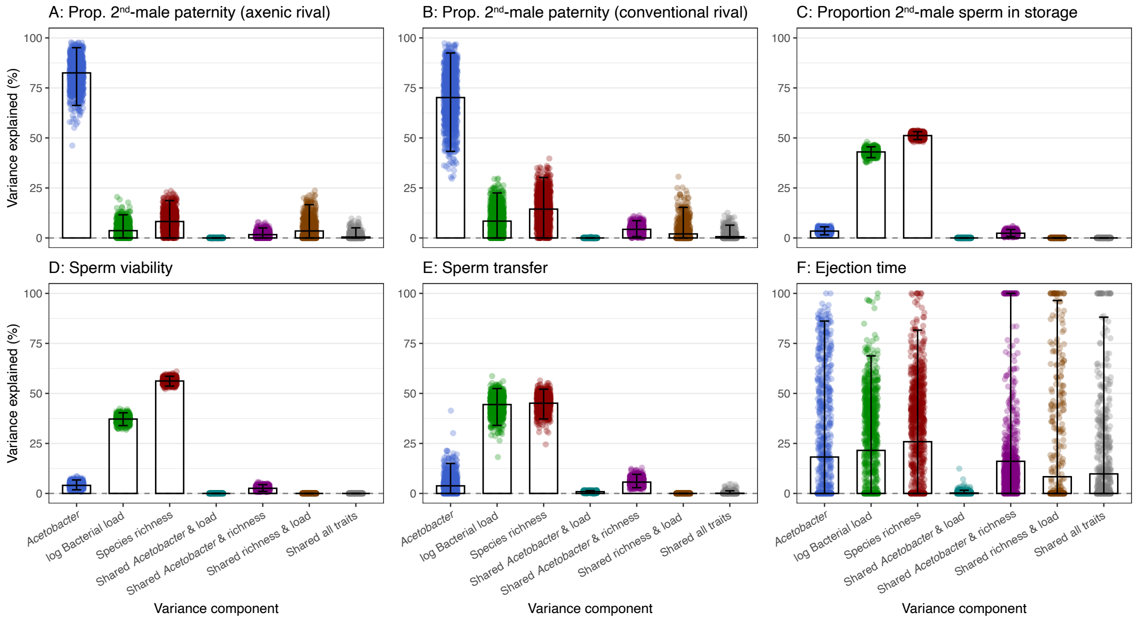


**Fig. S4:** Partitioning of clean variance components across reproductive traits. The figure shows the percentage of variance in each reproductive trait explained by unique and shared effects of relative *Acetobacter* abundance, species richness, and log bacterial load after renormalization ("clean" components). Each bar represents the mean ± 95% bootstrap confidence interval of the explained variance, and individual transparent points indicate bootstrap replicates. Shared components correspond to the variance jointly explained by multiple predictors.

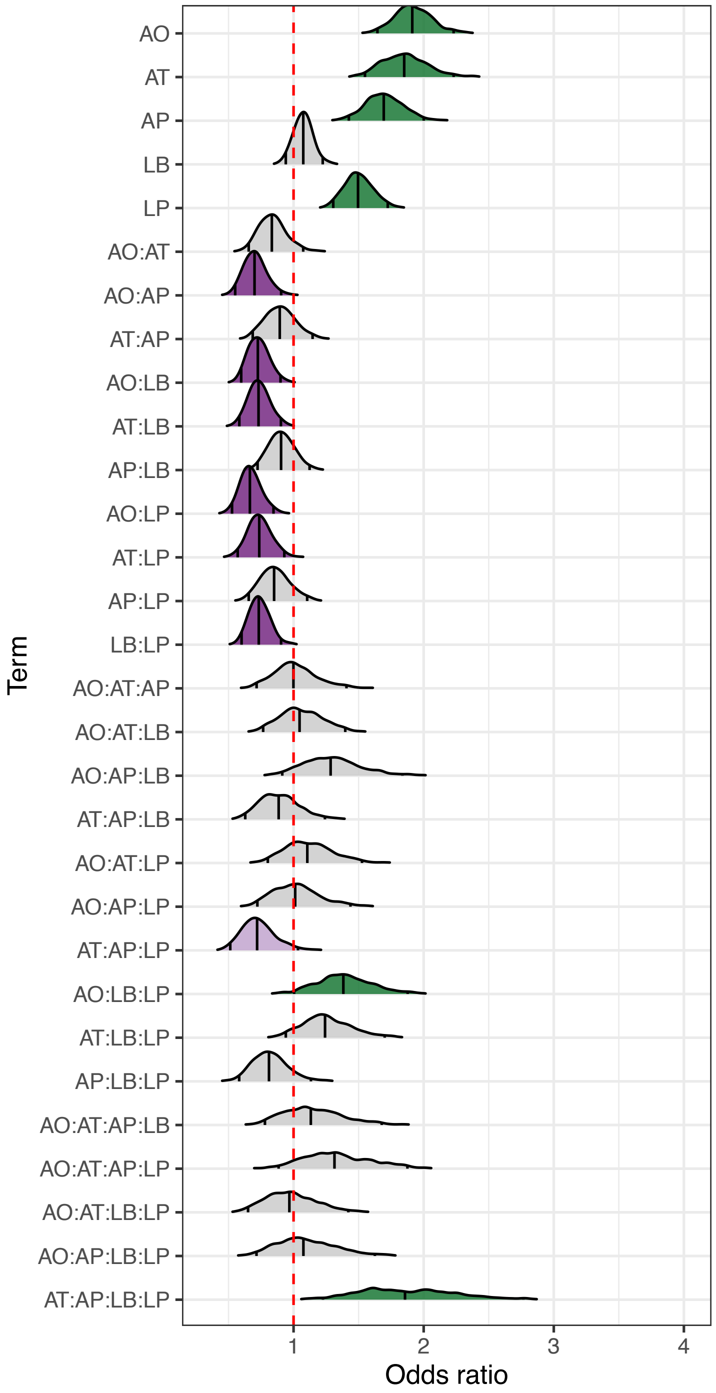


**Fig. S5.** Ridge plot showing the distribution of fixed-effect odds ratios across 1,000 bootstrap iterations of a generalized linear mixed-effects model on the proportion of focal-male progeny in sperm competition against axenic males. The model includes all main effects and two- to four-way interactions among five bacterial species. Each ridge represents the distribution of bootstrapped estimates for a given term, and its stability. Ridge colours indicate the direction of the effect (green: positive; purple: negative), with darker shades denoting effects whose 95% confidence intervals exclude zero, and lighter shades for those where only the 90% confidence interval excludes zero. The vertical dashed line indicates no effect.

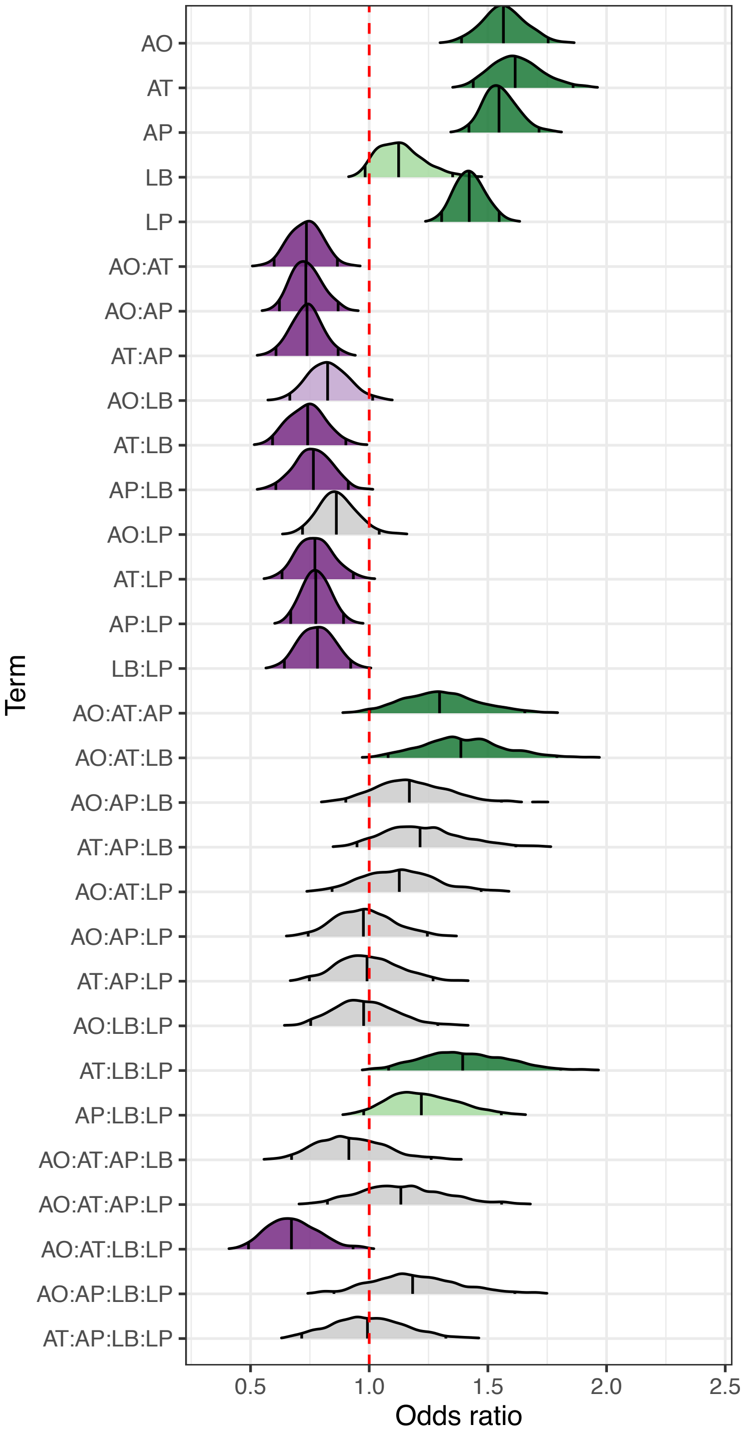


**Fig. S6.** Ridge plot showing the distribution of fixed-effect odds ratios across 1,000 bootstrap iterations of a generalized linear mixed-effects model on the proportion of focal-male progeny in sperm competition against males with an intact microbiome. The model includes all main effects and two- to four-way interactions among five bacterial species. Each ridge represents the distribution of bootstrapped estimates for a given term, and its stability. Ridge colours indicate the direction of the effect (green: positive; purple: negative), with darker shades denoting effects whose 95% confidence intervals exclude zero, and lighter shades for those where only the 90% confidence interval excludes zero. The vertical dashed line indicates no effect.

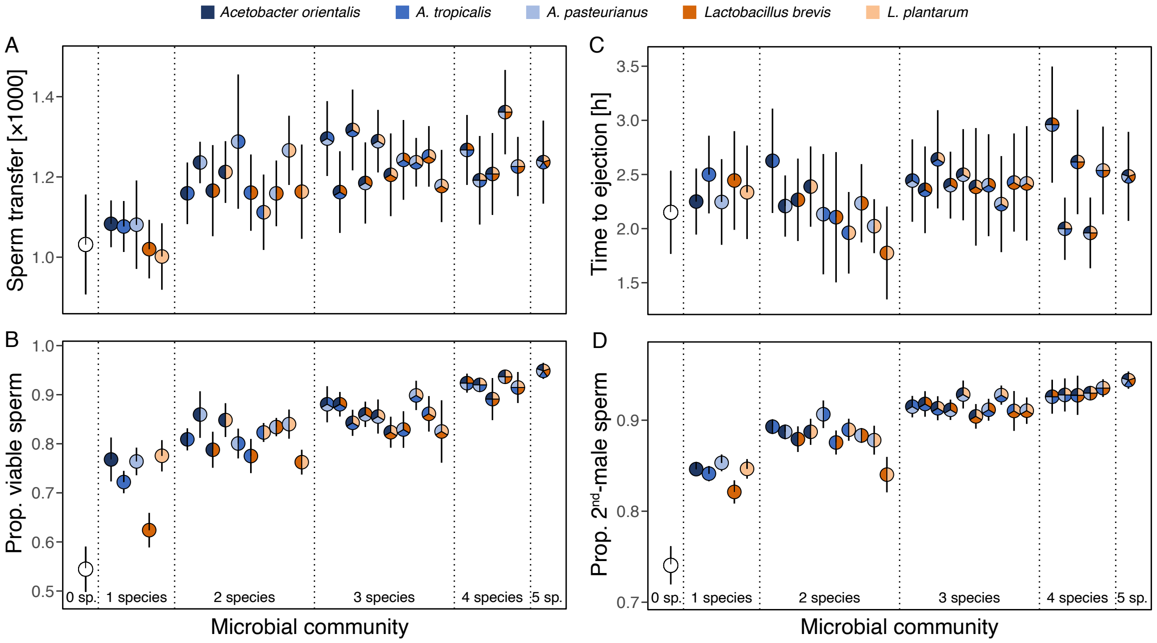


**Fig. S7.** Variation in male fitness traits relative to their microbial community. Colors indicate the different combinations of *Acetobacter* and *Lactobacillus* species associated with experimental males. Error bars in panel B reflect 95% confidence intervals from raw distributions (*N* = 20.2 ± 2.95 (± s.d.) trials per consortium).

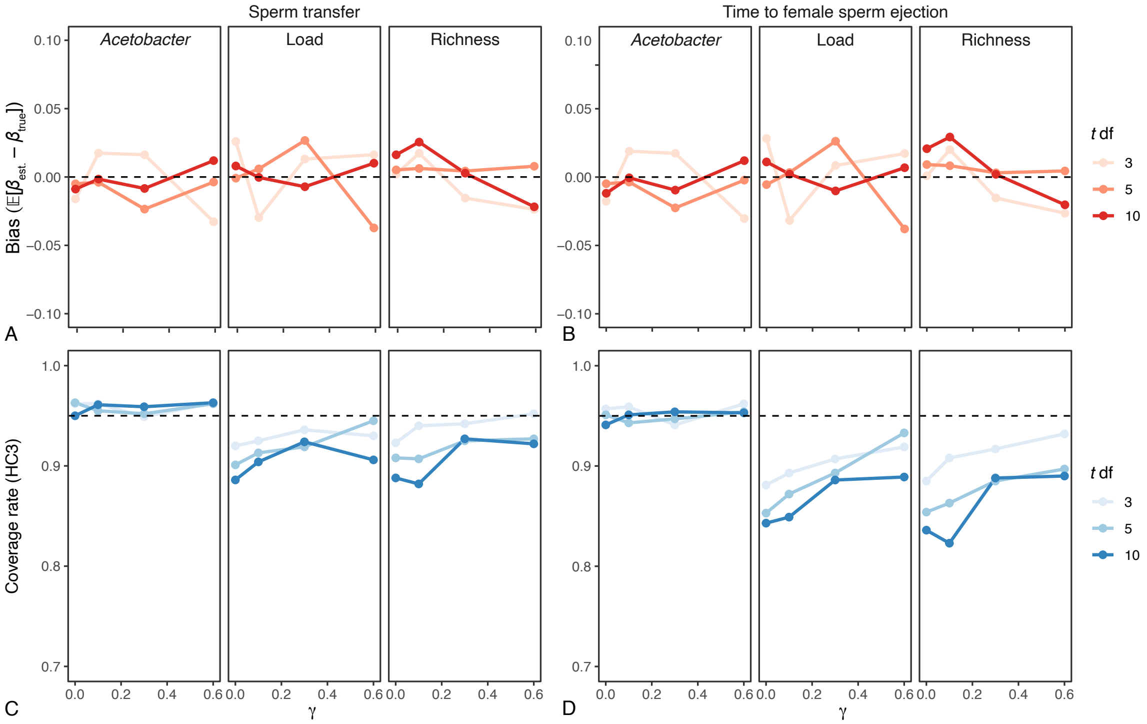


**Fig. S8.** Sensitivity of bias and 95%CI coverage to heteroskedasticity ($\gamma$) and tail heaviness ($t$ df) for models of sperm transfer (left) and ejection time (right). Each panel shows bias (top row; $\mathbb{E}[\beta_{estimated} - \beta_{true}]$ and coverage (bottom row; proportion of 95% CIs containing the true value) for three predictors under varying levels of heteroskedasticity ($\gamma$ = 0–0.6) and degrees of freedom of the $t$-distributed error term [$t$(df = 3, 5, 10)]. Increasing $\gamma$ introduces stronger variance inflation with predictor magnitude (i.e., greater heteroskedasticity), while lower $t$ df correspond to heavier-tailed error distributions (i.e., higher kurtosis).

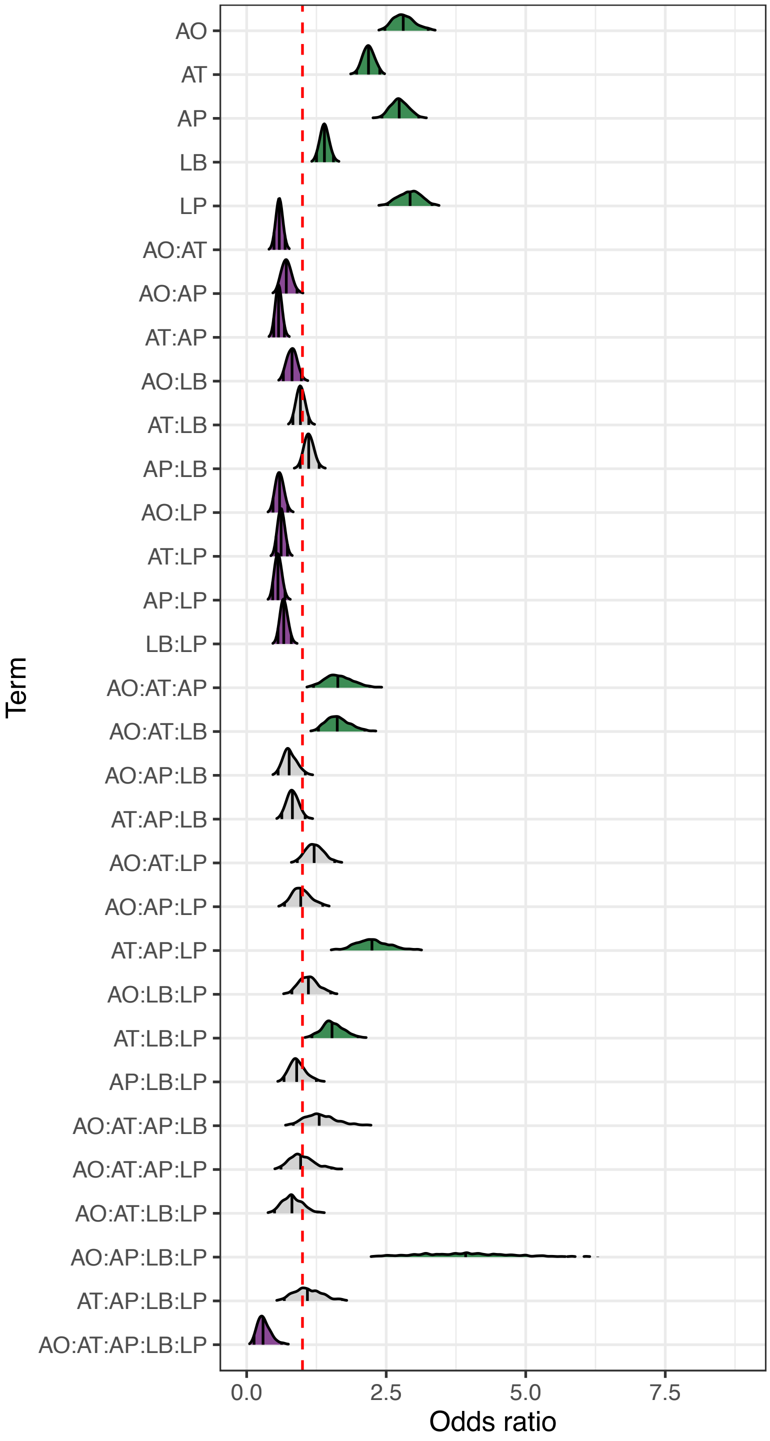


**Fig. S9.** Ridge plot showing the distribution of fixed-effect odds ratios across 1,000 bootstrap iterations of a generalized linear mixed-effects model on the proportion of viable sperm. The model includes all main effects and two- to four-way interactions among five bacterial species. Each ridge represents the distribution of bootstrapped estimates for a given term, and its stability. Ridge colours indicate the direction of the effect (green: positive; purple: negative) where 95% confidence intervals exclude zero. The vertical dashed line indicates no effect.

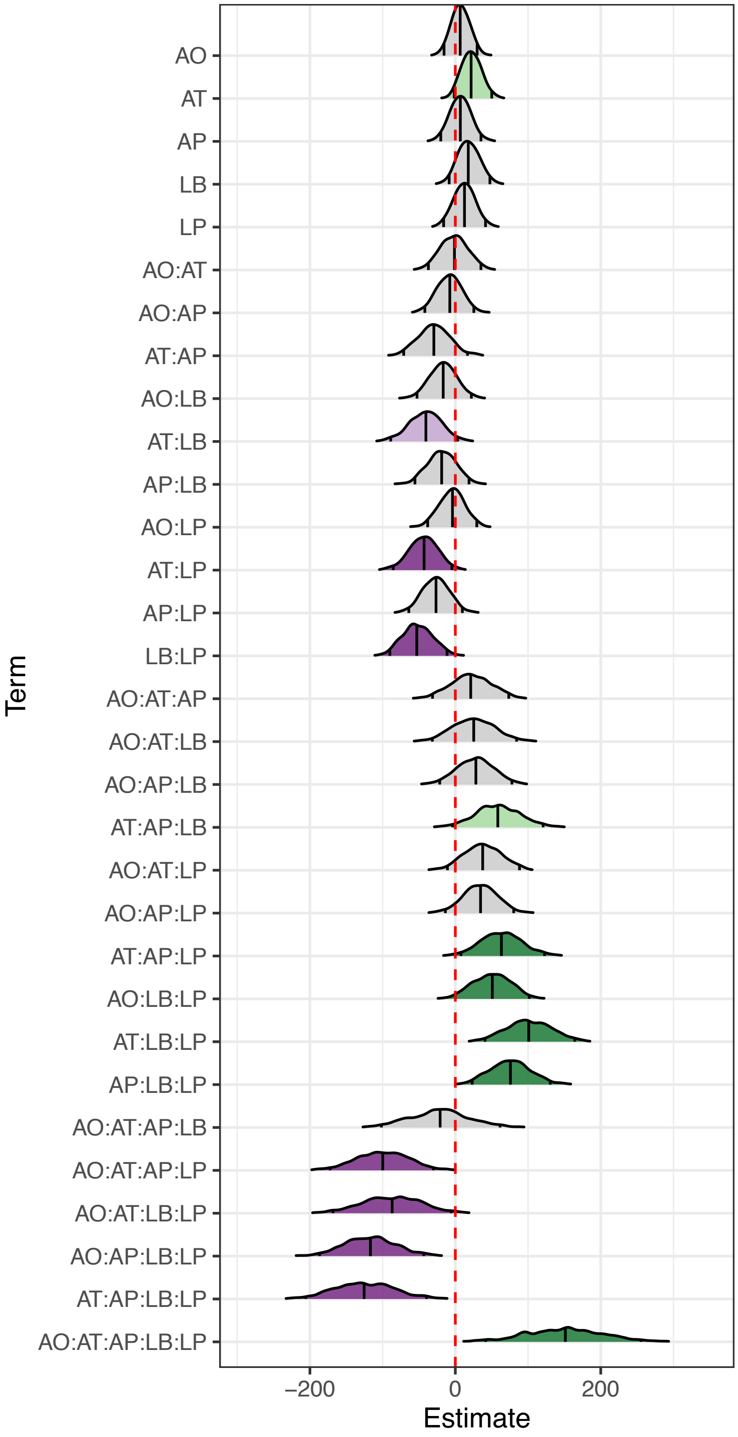


**Fig. S10.** Ridge plot showing the distribution of fixed-effect estimates across 1,000 bootstrap iterations of a linear mixed-effects model on the time to female sperm ejection. The model includes all main effects and two- to four-way interactions among five bacterial species. Each ridge represents the distribution of bootstrapped estimates for a given term, and its stability. Ridge colours indicate the direction of the effect (green: positive; purple: negative), with darker shades denoting effects whose 95% confidence intervals exclude zero, and lighter shades for those where only the 90% confidence interval excludes zero. The vertical dashed line indicates no effect. Estimates in minutes.

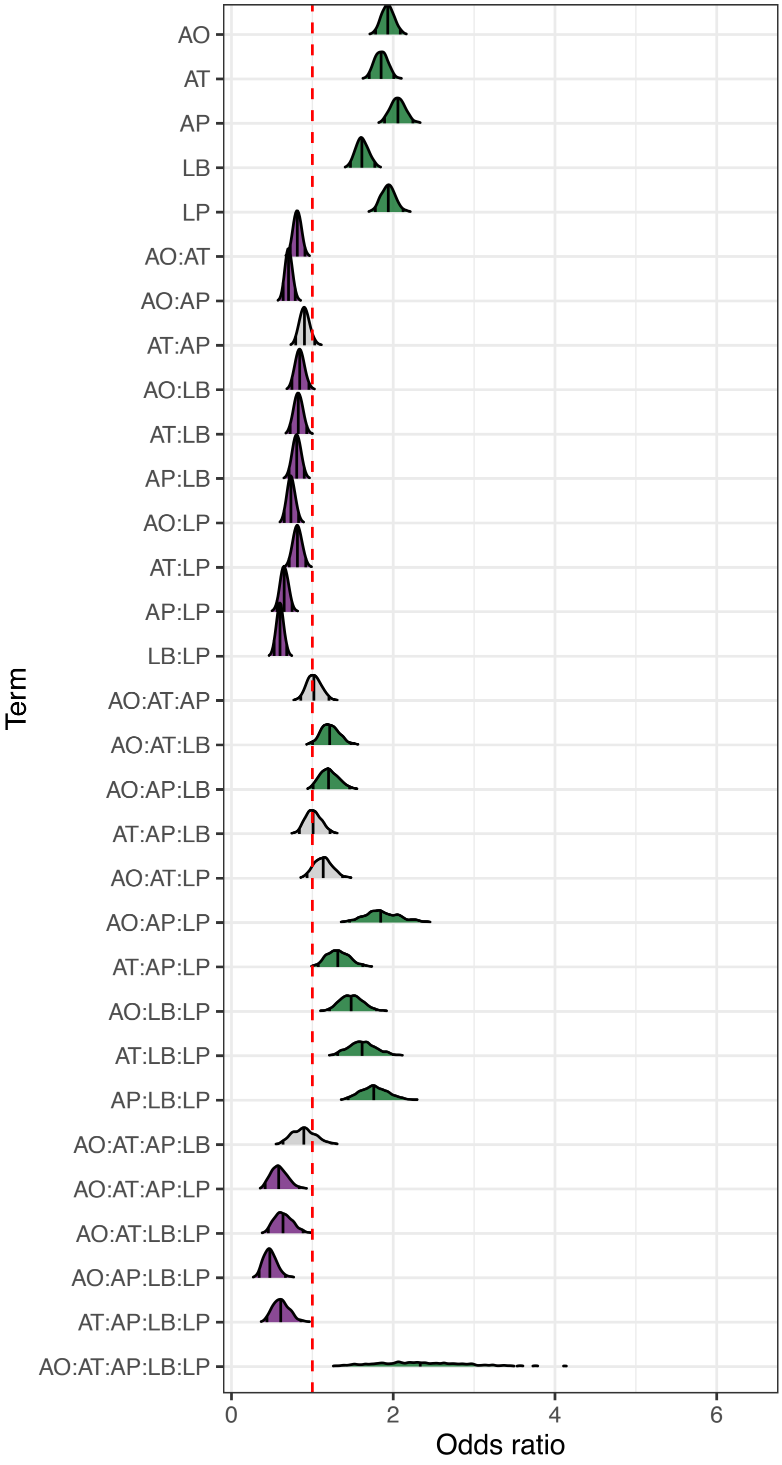


**Fig. S11.** Ridge plot showing the distribution of fixed-effect odds ratios across 1,000 bootstrap iterations of a generalized linear mixed-effects model on the proportion of focal-male sperm remaining in female storage after sperm ejection. The model includes all main effects and two- to four-way interactions among five bacterial species. Each ridge represents the distribution of bootstrapped estimates for a given term, and its stability. Ridge colours indicate the direction of the effect (green: positive; purple: negative) where 95% confidence intervals exclude zero. The vertical dashed line indicates no effect.

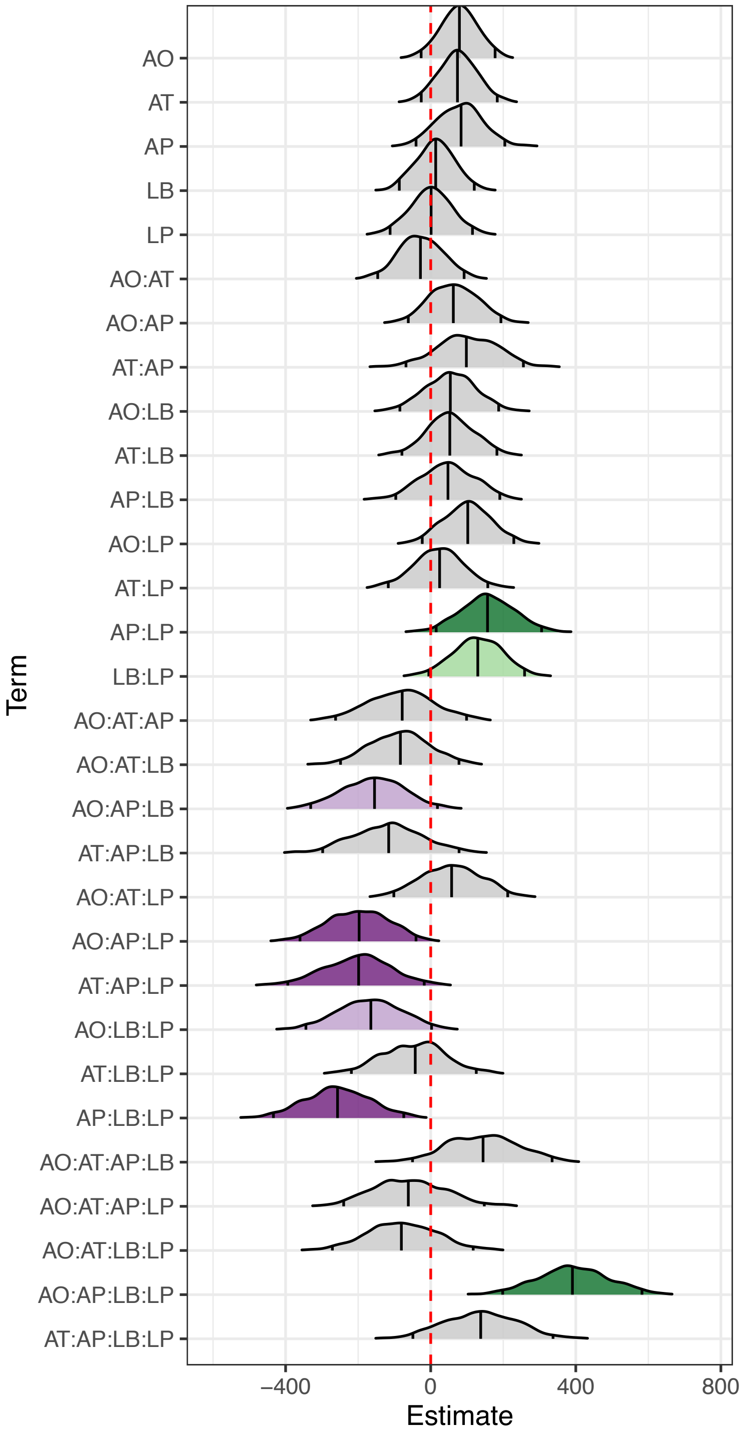


**Fig. S12.** Ridge plot showing the distribution of fixed-effect estimates across 1,000 bootstrap iterations of a linear mixed-effects model on the number of sperm transferred by focal males. The model includes all main effects and two- to four-way interactions among five bacterial species. Each ridge represents the distribution of bootstrapped estimates for a given term, and its stability. Ridge colours indicate the direction of the effect (green: positive; purple: negative), with darker shades denoting effects whose 95% confidence intervals exclude zero, and lighter shades for those where only the 90% confidence interval excludes zero. The vertical dashed line indicates no effect. Estimates in raw sperm counts.

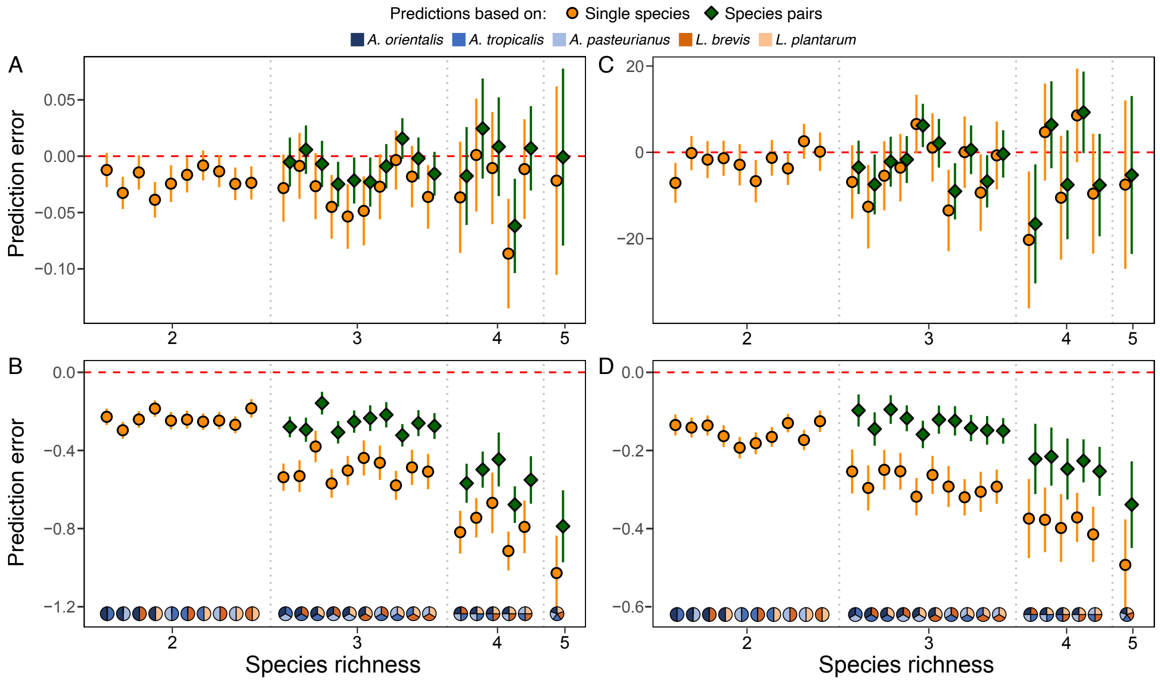


**Fig. S13.** Microbiome effects on (A) sperm transfer, (B) sperm viability, (C) time to female sperm ejection, and (D) the proportion of focal-male sperm stored by females. Each panel depicts the deviations from observed trait values when averaging those of the individual species (orange points) or species pairs (green diamonds) that jointly represented the same species communities (all coefficients derived from generalized linear mixed-effects models). Negative values indicate the observed trait value was higher than that predicted by averaging the effects of its component species (i.e., synergy), while a positive value indicates it was weaker (i.e., antagonism). The dashed horizontal line represents equality between the observed and the averaged effects. The *Acetobacter* and *Lactobacillus* species in each focal microbiome are indicated along the *x*-axis, and error bars reflect 95% confidence intervals from bootstrapped prediction differentials.

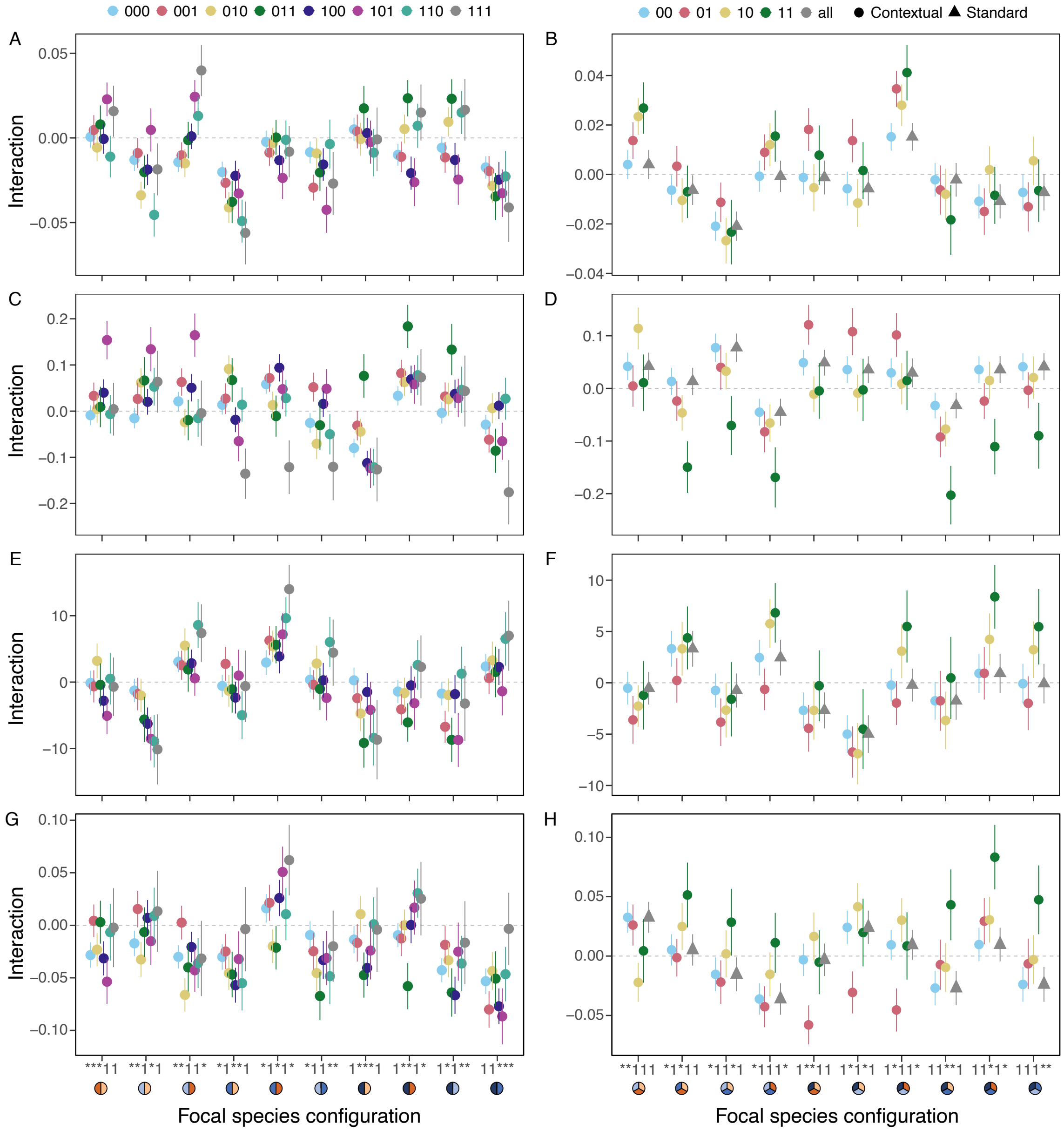


**Fig. S14.** Predictions of how given consortia of two (left) or three species affected (A, B) sperm transfer, (C, D) sperm viability, (E, F) the time to female sperm ejection, and (G, H) the proportion of focal-male sperm stored in the presence of varying bystander species. For each 5-digit code along the x-axes, 1's indicate the species configuration of the interaction (also visualized by pies underneath) and asterisks denote placeholders for bystander species (order within codes from left to right: *Acetobacter orientalis*, *A. tropicalis*, *A. pasteurianus*, *Lactobacillus brevis*, *L. plantarum*). The binary codes in the legend above each panel indicate the configuration of bystanders in the order, in which they would fill the placeholders (*) in each focal species configuration (1 = presence, 0 = absence). The right panels further distinguish between "standard" tests (grey triangles; i.e., global interaction across all community members present) and "contextual" tests (colored points; i.e., separate interactions between a focal species pair or trio for each bystander combination) per focal trio. All coefficients and propagated standard errors were derived from (generalized) linear mixed-effects models across 1,000 bootstrapped datasets.

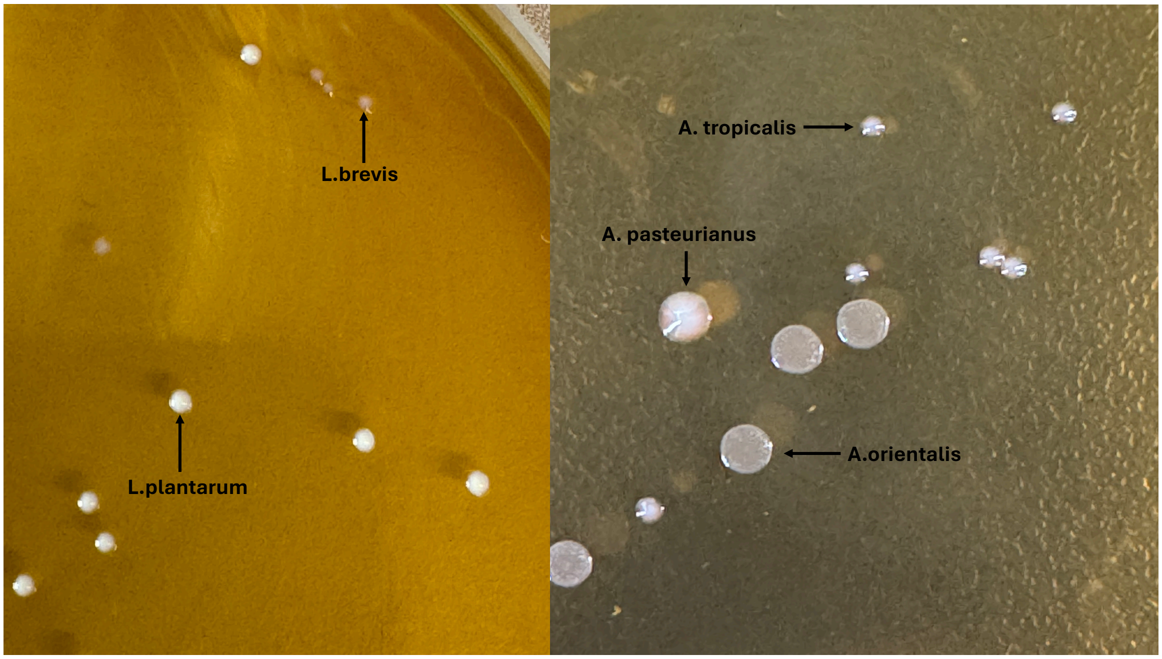


**Fig. S15.** Image showing morphological appearances of colonies formed by different bacterial species. On right side, acetobacter strains are present with *Acetobacter orientalis* represented by flat colonies with ruffled borders, *A. tropicalis* with small tan colored and highly reflective colonies, and *A. pasteurianus* with large tan colonies. In contrast on left side, *Lactobacillus brevis* shows brownish yellow colonies and *L. plantarum* with milky white colonies.

**Tables**

**Table S1.** Effects of different microbiome parameters on host fitness traits after residualizing species richness on bacterial load to address collinearity between these two variables. Effect sizes (with 95% confidence limits) are represented by partial correlation coefficients (*r*) for LMs and odds ratios (*OR*) for GLMs, along with their bootstrapped 95% confidence limits. Effects in bold highlight those that exclude 0 (for *r*) and 1 (for *OR*), respectively, from their confidence intervals. All variance inflation factors (VIF) were below the recommended threshold of 5 (e.g., O'Brien, 2007).

| **Response** | **Term** | ***r* [95% CL]** | ***OR* [95% CL]** | **VIF** |
|---|---|---|---|---|
| 2nd-male paternity | Proportion *Acetobacter* | | **1.21 [1.18, 1.25]** | 1.11 |
| (axenic rival) | log Bacterial load | | **1.05 [1.02, 1.08]** | 1.00 |
| | Species richness (resid) | | 1.02 [0.94, 1.11] | 1.11 |
| 2nd-male paternity | Proportion *Acetobacter* | | **1.10 [1.08, 1.13]** | 1.10 |
| (conventional rival) | log Bacterial load | | **1.04 [1.02, 1.06]** | 1.01 |
| | Proportion *Acetobacter* | | **1.08 [1.05, 1.10]** | 1.11 |
| 2nd-male sperm stored | Species richness (resid) | | 1.02 [0.95, 1.08] | 1.10 |
| | log Bacterial load | | **1.36 [1.32, 1.39]** | 1.02 |
| | Species richness (resid) | | **1.31 [1.21, 1.42]** | 1.11 |
| Sperm viability | Proportion *Acetobacter* | | **1.06 [1.02, 1.10]** | 1.14 |
| | log Bacterial load | | **1.49 [1.43, 1.54]** | 1.04 |
| | Species richness (resid) | | **1.98 [1.75, 2.22]** | 1.18 |
| Sperm transfer | Proportion *Acetobacter* | **0.33 [0.14, 0.52]** | | 1.09 |
| | log Bacterial load | **0.65 [0.51, 0.77]** | | 1.01 |
| | Species richness (resid) | 0.09 [-0.15, 0.32] | | 1.08 |
| Ejection time | Proportion *Acetobacter* | 0.17 [-0.07, 0.41] | | 1.09 |
| | log Bacterial load | **0.22 [0.00, 0.42]** | | 1.01 |
| | Species richness (resid) | 0.03 [-0.22, 0.28] | | 1.08 |

**Table S2.** Effects of different microbiome parameters on host fitness traits after residualizing bacterial load on species richness to address collinearity between these two variables. Effect sizes (with 95% confidence limits) are represented by partial correlation coefficients (*r*) for LMs and odds ratios (*OR*) for GLMs, along with their bootstrapped confidence limits. Effects in bold highlight those that exclude 0 (for *r*) and 1 (for *OR*), respectively, from their confidence intervals. All variance inflation factors (VIF) were below the recommended threshold of 5 (e.g., O'Brien, 2007).

| Response | Term | *r* [95% CL] | *OR* [95% CL] | VIF |
|---|---|---|---|---|
| $2^{nd}$-male paternity | Proportion *Acetobacter* | | **1.21 [1.18, 1.25]** | 1.11 |
| (axenic rival) | log Bacterial load (resid) | | 1.03 [0.94, 1.13] | 1.09 |
| | Species richness | | **1.05 [1.02, 1.08]** | 1.01 |
| $2^{nd}$-male paternity | Proportion *Acetobacter* | | **1.10 [1.08, 1.13]** | 1.10 |
| (conventional rival) | log Bacterial load (resid) | | 1.03 [0.96, 1.09] | 1.10 |
| | Species richness | | **1.04 [1.02, 1.06]** | 1.00 |
| $2^{nd}$-male sperm stored | Proportion *Acetobacter* | | **1.08 [1.05, 1.10]** | 1.11 |
| | log Bacterial load (resid) | | 1.06 [0.99, 1.13] | 1.12 |
| | Species richness | | **1.38 [1.34, 1.42]** | 1.02 |
| Sperm viability | Proportion *Acetobacter* | | **1.06 [1.02, 1.10]** | 1.14 |
| | log Bacterial load (resid) | | 0.80 [0.72, 0.89] | 1.18 |
| | Species richness | | **1.61 [1.54, 1.68]** | 1.05 |
| Sperm transfer | Proportion *Acetobacter* | **0.33 [0.14, 0.52]** | | 1.09 |
| | log Bacterial load (resid) | **0.24 [0.01, 0.46]** | | 1.09 |
| | Species richness | **0.63 [0.50, 0.76]** | | 1.00 |
| Ejection time | Proportion *Acetobacter* | 0.17 [-0.07, 0.41] | | 1.09 |
| | log Bacterial load (resid) | 0.06 [-0.17, 0.31] | | 1.09 |
| | Species richness | 0.21 [-0.01, 0.42] | | 1.00 |

**Table S3.** Effects of the proportional representation of *Acetobacter* and overall species richness on host fitness traits after restricting the dataset to observations within ± 1SD of the mean bacterial load in each bootstrap iteration (i.e., 23 of the 31 microbiome compositions). Effect sizes (with 95% confidence limits) are represented by partial correlation coefficients (*r*) for LMs and odds ratios (*OR*) for GLMs, along with their bootstrapped confidence limits. Effects in bold highlight those that exclude 0 (for *r*) and 1 (for *OR*), respectively, from their confidence intervals.

| Response | Term | *r* [95% CL] | *OR* [95% CL] |
|---|---|---|---|
| 2nd-male paternity | Proportion *Acetobacter* | | **1.31 [1.27, 1.36]** |
| (axenic rival) | Species richness | | 0.98 [0.93, 1.03] |
| 2nd-male paternity | Proportion *Acetobacter* | | **1.13 [1.09, 1.16]** |
| (conventional rival) | Species richness | | **1.04 [1.00, 1.09]** |
| 2nd-male sperm stored | Proportion *Acetobacter* | | **1.11 [1.07, 1.15]** |
| | Species richness | | **1.38 [1.32, 1.46]** |
| Sperm viability | Proportion *Acetobacter* | | **1.15 [1.10, 1.21]** |
| | Species richness | | **1.59 [1.47, 1.70]** |
| Sperm transfer | Proportion *Acetobacter* | **0.40 [0.15, 0.62]** | |
| | Species richness | **0.42 [0.14, 0.64]** | |
| Ejection time | Proportion *Acetobacter* | 0.16 [-0.16, 0.45] | |
| | Species richness | **0.38 [0.10, 0.63]** | |

**Table S4.** Effects of the proportional representation of *Acetobacter* and overall species richness on host fitness traits after excluding the bacterial load as a predictor. Effect sizes (with 95% confidence limits) are represented by partial correlation coefficients (*r*) for LMs and odds ratios (*OR*) for GLMs, along with their bootstrapped confidence limits. Effects in bold highlight those that exclude 0 (for *r*) and 1 (for *OR*), respectively, from their confidence intervals.

| **Response** | **Term** | ***r* [95% CL]** | ***OR* [95% CL]** |
|---|---|---|---|
| 2nd-male paternity | Proportion *Acetobacter* | | **1.21 [1.18, 1.24]** |
| (axenic rival) | Species richness | | **1.05 [1.02, 1.08]** |
| 2nd-male paternity | Proportion *Acetobacter* | | **1.10 [1.08, 1.12]** |
| (conventional rival) | Species richness | | **1.04 [1.02, 1.06]** |
| 2nd-male sperm stored | Proportion *Acetobacter* | | **1.07 [1.05, 1.09]** |
| | Species richness | | **1.39 [1.35, 1.43]** |
| Sperm viability | Proportion *Acetobacter* | | **1.09 [1.06, 1.13]** |
| | Species richness | | **1.58 [1.52, 1.64]** |
| Sperm transfer | Proportion *Acetobacter* | **0.28 [0.08, 0.46]** | |
| | Species richness | **0.62 [0.48, 0.74]** | |
| Ejection time | Proportion *Acetobacter* | 0.16 [-0.08, 0.40] | |
| | Species richness | 0.21 [-0.01, 0.41] | |

**Table S5.** Effects of the bacterial load and proportional representation of *Acetobacter* on host fitness traits after excluding species richness as a predictor. Effect sizes (with 95% confidence limits) are represented by partial correlation coefficients (*r*) for LMs and odds ratios (*OR*) for GLMs, along with their bootstrapped confidence limits. Effects in bold highlight those that exclude 0 (for *r*) and 1 (for *OR*), respectively, from their confidence intervals.

| **Response** | **Term** | ***r* [95% CL]** | ***OR* [95% CL]** |
|---|---|---|---|
| 2nd-male paternity | Proportion *Acetobacter* | | **1.22 [1.19, 1.25]** |
| (axenic rival) | log Bacterial load | | **1.05 [1.02, 1.08]** |
| 2nd-male paternity | Proportion *Acetobacter* | | **1.11 [1.08, 1.13]** |
| (conventional rival) | log Bacterial load | | **1.04 [1.02, 1.06]** |
| 2nd-male sperm stored | Proportion *Acetobacter* | | **1.11 [1.08, 1.14]** |
| | log Bacterial load | | **1.35 [1.32, 1.38]** |
| Sperm viability | Proportion *Acetobacter* | | **1.15 [1.11, 1.19]** |
| | log Bacterial load | | **1.44 [1.39, 1.48]** |
| Sperm transfer | Proportion *Acetobacter* | **0.36 [0.17, 0.54]** | |
| | log Bacterial load | **0.64 [0.51, 0.77]** | |
| Ejection time | Proportion *Acetobacter* | 0.18 [-0.06, 0.42] | |
| | log Bacterial load | **0.22 [0.00, 0.42]** | |

**Table S6.** Summary of regularized regression sensitivity analysis for microbiome predictors on host fitness traits. Shown are the mean, median, and standard deviation (SD) of coefficient estimates, along with 95% confidence limits across 1,000 bootstraps for each regularization type (α = 0 for ridge, 0.2 for elastic net, 1 for lasso). Predictors are standardized.

| Response | Term | α | Mean *β* | Median *β* | SD | 95% CL |
|---|---|---|---|---|---|---|
| 2nd-male paternity (axenic rival) | Prop. *Acetobacter* | 0 | 0.028 | 0.028 | 0.016 | [-0.01, 0.06] |
| | | 0.2 | 0.024 | 0.023 | 0.019 | [0.00, 0.06] |
| | | 1 | 0.018 | 0.009 | 0.022 | [0.00, 0.06] |
| | log Bacterial load | 0 | 0.029 | 0.027 | 0.018 | [0.00, 0.06] |
| | | 0.2 | 0.028 | 0.027 | 0.021 | [0.00, 0.07] |
| | | 1 | 0.028 | 0.026 | 0.026 | [0.00, 0.08] |
| | Species richness | 0 | **0.168** | **0.168** | **0.018** | **[0.13, 0.2]** |
| | | 0.2 | **0.174** | **0.176** | **0.018** | **[0.14, 0.21]** |
| | | 1 | **0.182** | **0.183** | **0.019** | **[0.14, 0.22]** |
| 2nd-male paternity (conventional rival) | Prop. *Acetobacter* | 0 | 0.018 | 0.018 | 0.014 | [-0.01, 0.04] |
| | | 0.2 | 0.017 | 0.016 | 0.016 | [0.00, 0.05] |
| | | 1 | 0.014 | 0.010 | 0.019 | [0.00, 0.05] |
| | log Bacterial load | 0 | 0.019 | 0.017 | 0.013 | [-0.01, 0.05] |
| | | 0.2 | 0.018 | 0.017 | 0.016 | [0.00, 0.05] |
| | | 1 | 0.019 | 0.018 | 0.019 | [0.00, 0.06] |
| | Species richness | 0 | **0.081** | **0.081** | **0.013** | **[0.06, 0.10]** |
| | | 0.2 | **0.084** | **0.085** | **0.012** | **[0.06, 0.11]** |
| | | 1 | **0.089** | **0.090** | **0.013** | **[0.06, 0.11]** |
| 2nd-male sperm stored | Prop. *Acetobacter* | 0 | **0.200** | **0.199** | **0.021** | **[0.16, 0.24]** |
| | | 0.2 | **0.233** | **0.230** | **0.040** | **[0.16, 0.32]** |
| | | 1 | **0.258** | **0.258** | **0.038** | **[0.19, 0.33]** |
| | log Bacterial load | 0 | **0.110** | **0.110** | **0.017** | **[0.08, 0.14]** |
| | | 0.2 | **0.084** | **0.089** | **0.032** | **[0.01, 0.14]** |
| | | 1 | 0.061 | 0.061 | 0.036 | [0.00, 0.13] |
| | Species richness | 0 | **0.070** | **0.069** | **0.011** | **[0.05, 0.09]** |
| | | 0.2 | **0.068** | **0.069** | **0.011** | **[0.05, 0.09]** |
| | | 1 | **0.064** | **0.065** | **0.014** | **[0.04, 0.09]** |
| Sperm viability | Prop. *Acetobacter* | 0 | **0.426** | **0.426** | **0.032** | **[0.36, 0.49]** |
| | | 0.2 | **0.620** | **0.628** | **0.082** | **[0.43, 0.76]** |
| | | 1 | **0.612** | **0.633** | **0.096** | **[0.42, 0.76]** |
| | log Bacterial load | 0 | 0.012 | 0.013 | 0.029 | [-0.05, 0.07] |
| | | 0.2 | -0.164 | -0.172 | 0.076 | [-0.29, 0.00] |
| | | 1 | -0.156 | -0.177 | 0.090 | [-0.29, 0.00] |
| | Species richness | 0 | **0.084** | **0.084** | **0.016** | **[0.05, 0.12]** |

| | | | | | | |
|---|---|---|---|---|---|---|
| | | 0.2 | **0.069** | **0.070** | **0.019** | **[0.03, 0.11]** |
| | | 1 | **0.066** | **0.067** | **0.019** | **[0.03, 0.10]** |
| Sperm transfer | Prop. *Acetobacter* | 0 | 0.233 | 0.241 | 0.100 | [-0.01, 0.42] |
| | | 0.2 | 0.209 | 0.228 | 0.148 | [-0.16, 0.46] |
| | | 1 | 0.197 | 0.180 | 0.187 | [0.00, 0.58] |
| | log Bacterial load | 0 | **0.331** | **0.299** | **0.125** | **[0.16, 0.62]** |
| | | 0.2 | **0.369** | **0.324** | **0.176** | **[0.12, 0.80]** |
| | | 1 | **0.406** | **0.436** | **0.208** | **[0.00, 0.73]** |
| | Species richness | 0 | **0.208** | **0.205** | **0.079** | **[0.07, 0.38]** |
| | | 0.2 | **0.214** | **0.215** | **0.092** | **[0.02, 0.39]** |
| | | 1 | **0.232** | **0.242** | **0.105** | **[0.00, 0.40]** |
| Ejection time | Prop. *Acetobacter* | 0 | 0.041 | 0.025 | 0.085 | [0.00, 0.19] |
| | | 0.2 | 0.030 | 0.000 | 0.101 | [0.00, 0.25] |
| | | 1 | 0.033 | 0.000 | 0.127 | [0.00, 0.33] |
| | log Bacterial load | 0 | 0.047 | 0.024 | 0.089 | [0.00, 0.21] |
| | | 0.2 | 0.046 | 0.000 | 0.114 | [0.00, 0.31] |
| | | 1 | 0.044 | 0.000 | 0.134 | [0.00, 0.36] |
| | Species richness | 0 | 0.052 | 0.015 | 0.078 | [-0.01, 0.27] |
| | | 0.2 | 0.050 | 0.000 | 0.092 | [0.00, 0.30] |
| | | 1 | 0.051 | 0.000 | 0.101 | [0.00, 0.35] |

**Table 7.** Summary of the percentage of variance in each reproductive trait explained by unique and shared effects of relative *Acetobacter* abundance, species richness, and log bacterial load, based on bootstrap variance partitioning. Values represent the mean percentages across bootstrap replicates, with 95% confidence limits derived from the bootstrap distributions, based on raw ("true") partitioning of variance (which may include negative contributions due to overlap among predictors), with shared components jointly explained by two or more predictors.

| Response | Component | Mean [95% CL] |
|---|---|---|
| 2nd-male paternity (axenic rival) | Proportion *Acetobacter* | 93.34 [81.34, 101.31] |
| | log Bacterial load | 4.15 [-1.64, 14.64] |
| | Species richness | 9.35 [-0.36, 23.60] |
| | Shared *Acetobacter* & load | 1.81 [-0.47, 5.56] |
| | Shared *Acetobacter* & richness | -3.77 [-6.63, -1.05] |
| | Shared richness & load | 1.16 [-14.29, 17.59] |
| | Shared all | -6.05 [-20.00, 5.11] |
| 2nd-male paternity (conventional rival) | Proportion *Acetobacter* | 82.73 [59.93, 99.92] |
| | log Bacterial load | 10.24 [-2.26, 30.16] |
| | Species richness | 17.73 [-0.77, 41.32] |
| | Shared *Acetobacter* & load | 5.14 [0.70, 11.66] |
| | Shared *Acetobacter* & richness | -2.75 [-5.38, -0.52] |
| | Shared richness & load | -5.69 [-30.26, 18.89] |
| | Shared all | -7.41 [-24.90, 8.18] |
| 2nd-male sperm stored | Proportion *Acetobacter* | 6.61 [3.08, 10.86] |
| | log Bacterial load | 79.34 [72.03, 86.06] |
| | Species richness | 94.50 [90.84, 97.44] |
| | Shared *Acetobacter* & load | 4.41 [1.46, 7.90] |
| | Shared *Acetobacter* & richness | -1.79 [-3.16, -0.63] |
| | Shared richness & load | -79.13 [-85.44, -71.29] |
| | Shared all | -3.94 [-8.03, -0.75] |
| Sperm viability | Proportion *Acetobacter* | 6.65 [3.11, 11.09] |
| | log Bacterial load | 60.93 [51.49, 69.14] |
| | Species richness | 91.69 [86.57, 95.37] |
| | Shared *Acetobacter* & load | 4.20 [1.73, 7.37] |
| | Shared *Acetobacter* & richness | -2.03 [-3.21, -1.03] |
| | Shared richness & load | -54.6 [-66.57, -40.54] |
| | Shared all | -6.84 [-11.14, -3.29] |
| Sperm transfer | Proportion *Acetobacter* | 6.80 [-7.34, 28.18] |
| | log Bacterial load | 84.53 [59.35, 104.15] |
| | Species richness | 86.39 [62.14, 106.91] |
| | Shared *Acetobacter* & load | 10.86 [5.56, 17.24] |
| | Shared *Acetobacter* & richness | 1.68 [0.20, 2.93] |
| | Shared richness & load | -84.97 [-105.58, -62.01] |
| | Shared all | -5.29 [-14.35, 2.25] |
| Ejection time | Proportion *Acetobacter* | 43.01 [-326.43, 465.73] |
| | log Bacterial load | 95.69 [-208.06, 729.99] |
| | Species richness | 148.79 [-116.50, 967.44] |
| | Shared *Acetobacter* & load | 42.6 [-0.98, 173.40] |
| | Shared *Acetobacter* & richness | -2.80 [-23.40, 7.86] |
| | Shared richness & load | -188.44 [-1210.08, 124.93] |
| | Shared all | -38.86 [-374.97, 93.82] |

**Table S8.** Renormalized variance components (with 95% confidence limits) from Table S7 such that all contributions are positive and sum to 100% of the total explained variance (i.e., “clean” variance components), facilitating comparison of relative effect sizes. These values correspond to those shown in Fig. S4.

| Response | Component | Mean [95% CL] |
|---|---|---|
| 2nd-male paternity (axenic rival) | Proportion *Acetobacter* | 82.88 [65.28, 95.47] |
| | log Bacterial load | 3.70 [0.00, 11.97] |
| | Species richness | 8.15 [0.00, 19.09] |
| | Shared *Acetobacter* & load | 1.61 [0.00, 4.72] |
| | Shared *Acetobacter* & richness | 0.00 [0.00, 0.00] |
| | Shared richness & load | 3.21 [0.00, 14.26] |
| | Shared all | 0.45 [0.00, 4.66] |
| 2nd-male paternity (conventional rival) | Proportion *Acetobacter* | 70.14 [43.20, 92.52] |
| | log Bacterial load | 8.36 [0.00, 21.56] |
| | Species richness | 14.42 [0.00, 30.01] |
| | Shared *Acetobacter* & load | 4.24 [0.55, 9.11] |
| | Shared *Acetobacter* & richness | 0.00 [0.00 0.00] |
| | Shared richness & load | 2.16 [0.00, 14.72] |
| | Shared all | 0.68 [0.00, 6.68] |
| 2nd-male sperm stored | Proportion *Acetobacter* | 3.58 [1.65, 5.97] |
| | log Bacterial load | 42.91 [39.79, 45.7] |
| | Species richness | 51.13 [49.21, 53.23] |
| | Shared *Acetobacter* & load | 2.39 [0.78, 4.32] |
| | Shared *Acetobacter* & richness | 0.00 [0.00, 0.00] |
| | Shared richness & load | 0.00 [0.00, 0.00] |
| | Shared all | 0.00 [0.00, 0.00] |
| Sperm viability | Proportion *Acetobacter* | 4.08 [1.92, 6.96] |
| | log Bacterial load | 37.23 [33.89, 40.09] |
| | Species richness | 56.13 [53.76, 58.58] |
| | Shared *Acetobacter* & load | 2.57 [1.07, 4.48] |
| | Shared *Acetobacter* & richness | 0.00 [0.00, 0.00] |
| | Shared richness & load | 0.00 [0.00, 0.00] |
| | Shared all | 0.00 [0.00, 0.00] |
| Sperm transfer | Proportion *Acetobacter* | 4.29 [0.00, 16.35] |
| | log Bacterial load | 44.06 [34.19, 52.44] |
| | Species richness | 44.96 [36.96, 51.62] |
| | Shared *Acetobacter* & load | 5.74 [2.76, 9.38] |
| | Shared *Acetobacter* & richness | 0.87 [0.11, 1.45] |
| | Shared richness & load | 0.00 [0.00, 0.00] |
| | Shared all | 0.08 [0.00, 1.32] |
| Ejection time | Proportion *Acetobacter* | 18.99 [0.00, 85.55] |
| | log Bacterial load | 22.23 [0.00, 66.56] |
| | Species richness | 26.93 [0.00, 79.71] |
| | Shared *Acetobacter* & load | 14.7 [0.00, 100.00] |
| | Shared *Acetobacter* & richness | 0.23 [0.00, 1.66] |
| | Shared richness & load | 8.21 [0.00, 93.34] |
| | Shared all | 8.71 [0.00, 81.37] |

**Table S9:** Results of forward-simulation validation assessing bias, variance, mean squared error (MSE), and 95% confidence-interval (CI) coverage for each predictor in the fitness models. For each response variable, 1,000 datasets were simulated using the empirically estimated coefficients as the true generative parameters, with predictors drawn from a multivariate normal distribution preserving the observed correlation between bacterial load and species richness ($\rho$ = 0.51). Models were refit to each simulated dataset using the same structure as the empirical analysis, and inference was obtained using HC3 heteroskedasticity-consistent confidence intervals. For each predictor, *True β* denotes the generative effect size; *Mean estimate* is the average recovered coefficient across simulations; *Bias* is the mean error ($\mathbb{E}[\beta_{estimated} - \beta_{true}]$); *MSE* is the mean squared error ($\mathbb{E}[\beta_{estimated} - \beta_{true}]^2$); and *Coverage rate* is the proportion of simulations in which the 95% HC3 CI contained the true $\beta$.

| Response | Predictor | True *β* | Mean est. | Bias | MSE | Coverage rate |
|---|---|---|---|---|---|---|
| 2nd-male paternity | Prop. *Acetobacter* | 0.190 | 0.204 | 0.014 | 0.370 | 0.990 |
| (axenic rival) | log Bacterial load | 0.030 | 0.069 | 0.039 | 0.381 | 0.991 |
| | Species richness | 0.020 | 0.036 | 0.016 | 0.361 | 0.988 |
| 2nd-male paternity | Prop. *Acetobacter* | 0.100 | 0.088 | -0.012 | 0.367 | 0.983 |
| (conventional rival) | log Bacterial load | 0.020 | 0.045 | 0.025 | 0.385 | 0.988 |
| | Species richness | 0.020 | 0.040 | 0.020 | 0.356 | 0.992 |
| 2nd-male sperm stored | Prop. *Acetobacter* | 0.070 | 0.046 | -0.024 | 0.349 | 0.991 |
| | log Bacterial load | 0.060 | 0.094 | 0.034 | 0.398 | 0.983 |
| | Species richness | 0.270 | 0.351 | 0.081 | 0.414 | 0.977 |
| Sperm viability | Prop. *Acetobacter* | 0.060 | 0.049 | -0.011 | 0.336 | 0.990 |
| | log Bacterial load | -0.230 | -0.245 | -0.015 | 0.418 | 0.983 |
| | Species richness | 0.680 | 0.853 | 0.173 | 0.523 | 0.986 |
| Sperm transfer | Prop. *Acetobacter* | 0.260 | 0.253 | -0.007 | 0.060 | 0.944 |
| | log Bacterial load | 0.470 | 0.485 | 0.015 | 0.121 | 0.850 |
| | Species richness | 0.170 | 0.185 | 0.015 | 0.111 | 0.868 |
| Ejection time | Prop. *Acetobacter* | 0.170 | 0.162 | -0.008 | 0.068 | 0.929 |
| | log Bacterial load | 0.160 | 0.179 | 0.019 | 0.163 | 0.781 |
| | Species richness | 0.060 | 0.078 | 0.018 | 0.155 | 0.808 |
